# Psilocybin Attenuates Cortical Representations of Aversion in the Mouse Auditory Cortex

**DOI:** 10.64898/2026.03.26.714498

**Authors:** James D. Johnson, Runyi Tian, Yasaman Etemadi, Zheng Li

## Abstract

Psilocybin can produce sustained benefits in affective and trauma-related disorders, yet if and how it reshapes sensory representations of learned valence associations remains largely unclear. To address this, we used longitudinal two-photon calcium imaging in awake C57BL/6 mice to examine how psilocybin modulates layer 2/3 auditory cortex activity at single-cell and population levels. Evoked responses were measured for tones with or without prior associations with valenced stimuli, as well as for the valenced stimuli themselves. Most responsive neurons were selective for tones alone, while distinct subsets responded exclusively to reward or aversive stimuli, and a smaller population encoded both. Psilocybin selectively reduced responses to aversive stimuli and earlier-established aversive-associated tones, without affecting aversive association, reward responses, or responses to newly aversive-associated tones. At the population level, psilocybin acutely increased coordination across tone-responsive neurons, while later reducing it selectively among neurons encoding the aversive-associated tone. These results demonstrate that psilocybin preferentially dampens well consolidated aversive sensory representations in auditory cortex, rather than fresh associations, without broadly affecting auditory processing or new aversive learning.

## Introduction

Psilocybin is a classic serotonergic psychedelic that alters perception, mood, and cognition, with effects mediated in part by agonism at the 5-HT₂A receptor, while also engaging other serotonin receptor subtypes and non-serotonergic mechanisms [1,2]. In preclinical studies, psilocybin has been associated with increased neuroplasticity, including dendritic spine formation, synaptogenesis, and glutamatergic receptor changes [3-5]. These effects may contribute to durable reorganization of neural circuitry. Consistent with this, both animal studies and clinical trials suggest therapeutic potential across several conditions, including major depressive disorder (MDD), PTSD, addiction, and pain [6-13]. Notably, physiological and behavioral benefits can outlast psilocybin’s pharmacokinetics, persisting for weeks to months after a single dose [3,14]. Human neuroimaging studies indicate that psilocybin modulates large-scale functional networks, including acute and longer-lasting alterations in default mode network connectivity, which may create windows for restructuring maladaptive patterns of activity [15-17]. In this framework, psilocybin is often proposed to relax rigid negative affective states, but reported effects on fear extinction and aversive learning have been variable across studies [8,11,18-20]. Such variability likely reflects differences in circuit engagement, memory strength, and timing relative to consolidation [21].

Consistent with this view, serotonergic receptors are broadly expressed across the brain, and psilocybin’s effects appear to depend on region- and cell-type-specific mechanisms. At the circuit level, long-term behavioral effects can depend on engagement of specific neuronal subpopulations within association cortices such as prefrontal cortex [14]. In the retrosplenial cortex, psilocybin has been reported to recruit previously inactive neuronal populations while suppressing fear-active ensembles and enhancing fear extinction; a process which may also be partially mediated by psilocybin through hippocampal plasticity [11,19]. Despite promising clinical outcomes under controlled conditions, the mechanisms through which psilocybin alters responses to negatively valenced cues, and how prior aversive associations or newly encountered emotionally salient experiences are updated during and after dosing, remain incompletely understood. This gap is particularly relevant for understanding how sensory reminders of emotionally salient experiences are processed and potentially modified.

The auditory cortex provides a useful model system for addressing these questions. It plays a central role in processing sound features such as frequency and timing and exhibits substantial experience-dependent plasticity [22]. Importantly, sensory cortices are increasingly recognized as sites of associative and emotional integration rather than passive feature detectors, receiving valence-related inputs from limbic and association regions such as the amygdala [23,24]. Aberrant auditory processing is implicated in disorders such as schizophrenia and auditory verbal hallucinations, and altered auditory perception is commonly reported during psychedelic experiences. Auditory cortical responses can be shaped by non-auditory information, including valence and contextual signals. In humans, operant and classical conditioning modulate auditory cortex responses during task performance [25], pleasant and unpleasant sounds led to increased auditory cortex activation as compared to neutral sounds [26], and auditory cortex has been shown to encode non-auditory variables such as behavioral relevance and context [27]. In animal models, pairing an auditory conditioned stimulus with foot shock can induce auditory cortical responses to the aversive stimulus alone, indicating the formation of novel sensory-valence associations [28]. Together with broader evidence that sensory cortices participate in emotional processing and valence integration [29-34], these findings suggest that auditory cortex is a downstream target of affective modulation. While 5-HT₂A receptors are expressed in the auditory cortex [35], it is not clear if the effects of psilocybin arise from direct local actions or through modulation of interconnected sensory association and limbic regions.

While neuroimaging and circuit-mapping studies of psilocybin have focused primarily on frontal, association and limbic regions, sensory cortices have received comparatively less attention [5,36,37]. Across the cortex, 5-HT₂A agonist effects appear heterogeneous; association cortices such as ACC show net increases in spiking with altered synchrony, whereas primary sensory cortices show context- and circuit-dependent changes in response gain and variability, consistent with findings in visual cortex suggesting that psychedelic modulation reflects stimulus context rather than uniform excitation or suppression [38-40]. In the auditory cortex, two-photon imaging studies have reported an immediate transient hyperexcitability followed by response suppression and altered habituation without tonotopic reorganization or alterations in acoustic processing [41,42]. However, how psilocybin affects auditory cortical processing under emotionally salient conditions, as encountered in psilocybin-assisted psychotherapy, remains unclear.

Here, we used longitudinal two-photon calcium imaging of auditory cortex neurons in awake mice to examine how psilocybin modulates sensory cortical responses to emotionally valenced stimuli. We first characterized neural selectivity for tones, reward, and aversive stimuli, presented both in paired and unpaired contexts. We then focused on negative valence, tracking auditory cortical representations of established and newly acquired aversive associations across three weeks following psilocybin administration. By repeatedly imaging the same neurons over time, we reveal how psilocybin shapes the encoding of aversive experience at the level of individual cells and subpopulations in sensory cortex.

## Materials and Methods

### Animals

All procedures conformed to NIH guidelines and were approved by the National Institute of Mental Health Animal Care and Use Committee. A total of 33 C57BL/6J mice (Charles River Laboratories) were used and maintained on a 12-h light/dark cycle. During imaging sessions involving water, mice had ad libitum access to food and were water restricted to 15 min of water access per day, in addition to water obtained during experiments. Body weight was monitored before each session to ensure maintenance of at least 85% of baseline pre-experiment weight. Mice with poor imaging quality, including cloudy cranial windows, weak GCaMP expression, unstable signal-to-noise ratio, or absent or weak tone responses, were excluded.

### Cranial window implantation and AAV injection

Mice received intraperitoneal xylazine (5 mg/kg) and dexamethasone (30 mg/kg, # 2392-39-4) for perioperative analgesia and anti-inflammatory support, then were anesthetized with isoflurane (5% induction, 1-1.5% maintenance) in a stereotaxic frame. Ophthalmic ointment (Dechra, Puralube) was applied, and body temperature was maintained at 37.5 °C. The scalp was shaved, disinfected with iodine, and incised to expose the skull, which was cleaned with artificial cerebrospinal fluid (aCSF), and the skin edges were sealed with Cyanoacrylate adhesive (Vetbond, #1469SB). A craniotomy was performed over the right auditory cortex (AP: 2.45-2.6 mm, ML: 4.35-4.5 mm, DV: -0.3 to -0.5 mm), and the dura was kept moist with aCSF. AAV-syn-GCaMP8f (Addgene, #162376; 5×10¹²–1×10¹³ GC/mL) was injected (300 nL per site, 90 nL per minute) into 4-5 posterior sites spaced ∼0.5 mm apart to span estimated primary auditory cortex. Double coverslips were prepared for cranial windows consisting of a 3 mm bonded (Norland, NOA 71) to 4 mm round coverslip (Warner, #64-0720, #64-0724). Double coverslips were affixed over the craniotomy using cyanoacrylate adhesive (Krazy Glue, #KG58548R) and further sealed dental cement. A Y-shaped titanium head bar was cemented to the contralateral skull, and mice were singly housed post-operatively to prevent window damage. Post-operative care consisted of 0.1ml Ketoprofen for 3 days post-surgery.

### Two-Photon Imaging

Imaging was performed on a Bruker Ultima two-photon two-photon microscope with a custom head-fixation stage. Two-photon excitation was delivered at 910 nm (Chameleon Vision S, Coherent) at 20-30 mW at the sample. After 3-4 weeks of recovery, mice were habituated to head fixation on a floating Styrofoam ball (PhenoSys JetBall TFT System. PhenoSys GmbH) for at least 2 h across multiple sessions and trained to lick from a motorized waterspout. Layer 2/3 neurons (100-250 µm below pia) were imaged using a 25× water-immersion objective (NA 1.05, Olympus, XLPLN25XWMP2) mounted on an orbital nosepiece (35-50° tilt). Imaging data were acquired using resonant galvo meters at approximately 30 Hz (512 × 512 pixels). A 60:40 ultrasound gel to water mix was used as the immersion medium to allow for chronic imaging without evaporation (Aquasonic Clear, #03-50, refractive index ∼1.33-1.35). Images were acquired using GaAsP photomultiplier tubes via Prairie View software (version 5.4, Bruker Corporation). The same field of view (∼400 × 400 µm) and neurons were re-identified and imaged across sessions using saved stage coordinates, surface vasculature landmarks, and alignment to the first sessions mean or max projection image. Power was kept as close as possible between sessions for each mouse, and signal to noise (SNR) was calculated across all images to ensure maintenance of image quality, with no differences found in power or SNR across groups or sessions (Supplementary Figure 1). All mice were imaged while awake, head fixed, on an air-cushioned spherical treadmill, with a plastic tube kept above them to aid stability.

### Auditory and valence stimulation

Auditory stimuli were generated in MATLAB (MathWorks, R2025b) and delivered using an Avisoft SASLab Pro/Recorder software and UltraSoundGate Player 116H (Avisoft Bioacoustics, Inc.); a free-standing speaker positioned 10 cm lateral to the ear contralateral to the cranial window. Across sessions, all tones had 10 ms cosine ramps and were presented at ∼70 dB SPL measured at the mouse head position with a calibrated SPL meter. After habituation, this level did not elicit overt behavioural responses. For tonotopic mapping, mice were presented with pseudorandom pure tones spanning 4 to 35 kHz (5 tones, approximately 20 repeats each, 1 s duration, 10 s inter-tone interval). Imaging fields of view for subsequent valence experiments were selected from regions showing robust and consistent tone responsiveness within the 4 to 25 kHz range. For all tone blocks the utilized tones were generated and presented in the same pipeline as above. Valence stimuli were delivered with PhenoSys hardware in separate blocks from tones. Water (5 µL per trial) was delivered via a motorized lick port with a lick sensor at the tip to confirm successful licks. Air-puff delivery consisted of a metal puff tube and connected via plastic tubing to a valve, which was supplied by a compressed-airline regulated through a flow meter and pressure regulator. Air puffs consisted of a 0.5 s, 15-PSI air puff directed at the ipsilateral eye relative to the imaging site, delivered from 10 cm. To minimize auditory artifacts, the valve was relocated away from the imaging stage and mechanically damped. Water and air-puff delivery did not produce detectable sound with a calibrated SPL meter above baseline at the mouse’s head position.

### Reward and aversive dataset paradigm (Figure 1)

On Day 1 (pairing), mice were presented with a tone block containing three tones separated by at least half an octave (4, 8, and 13 kHz). The 8 kHz and 13 kHz tones were paired by co-termination with either water or air puff (counterbalanced across mice), and the 4 kHz tone remained unpaired. Each trial consisted of a 1 s tone and, when paired, a 0.5 s stimulus that co-terminated with the tone. Trials were separated by a 11 to 17 s pseudorandom inter-trial interval, with around 30 repeats per tone. On Day 2, the same auditory cortex field of view was re-identified and re-imaged. Each imaging session consisted of a dedicated stimulus block, and a dedicated tone block. The stimulus block presented water trials, air puff trials, and non-evoked trials with no tone or stimulus (partial lick port extension without water delivery) in pseudorandom order to ensure around 20 presentations of each stimulus. The tone block presented pseudo randomly pure tones (4, 8, and 13 kHz; 1 s) with an equal number of repeats per frequency (∼20 presentations each). Mice were imaged in an ∼11-minute Pre and Post session, before and after intraperitoneal saline or psilocybin, in awake mice (2 mg/kg in 0.1 mL saline, Cayman: #14041). Post imaging blocks began 10 minutes after injection and were completed within 35 minutes of injection. On injection days, topical lidocaine was applied to the shaved abdomen 30 minutes before intraperitoneal injection.

**Figure 1.**
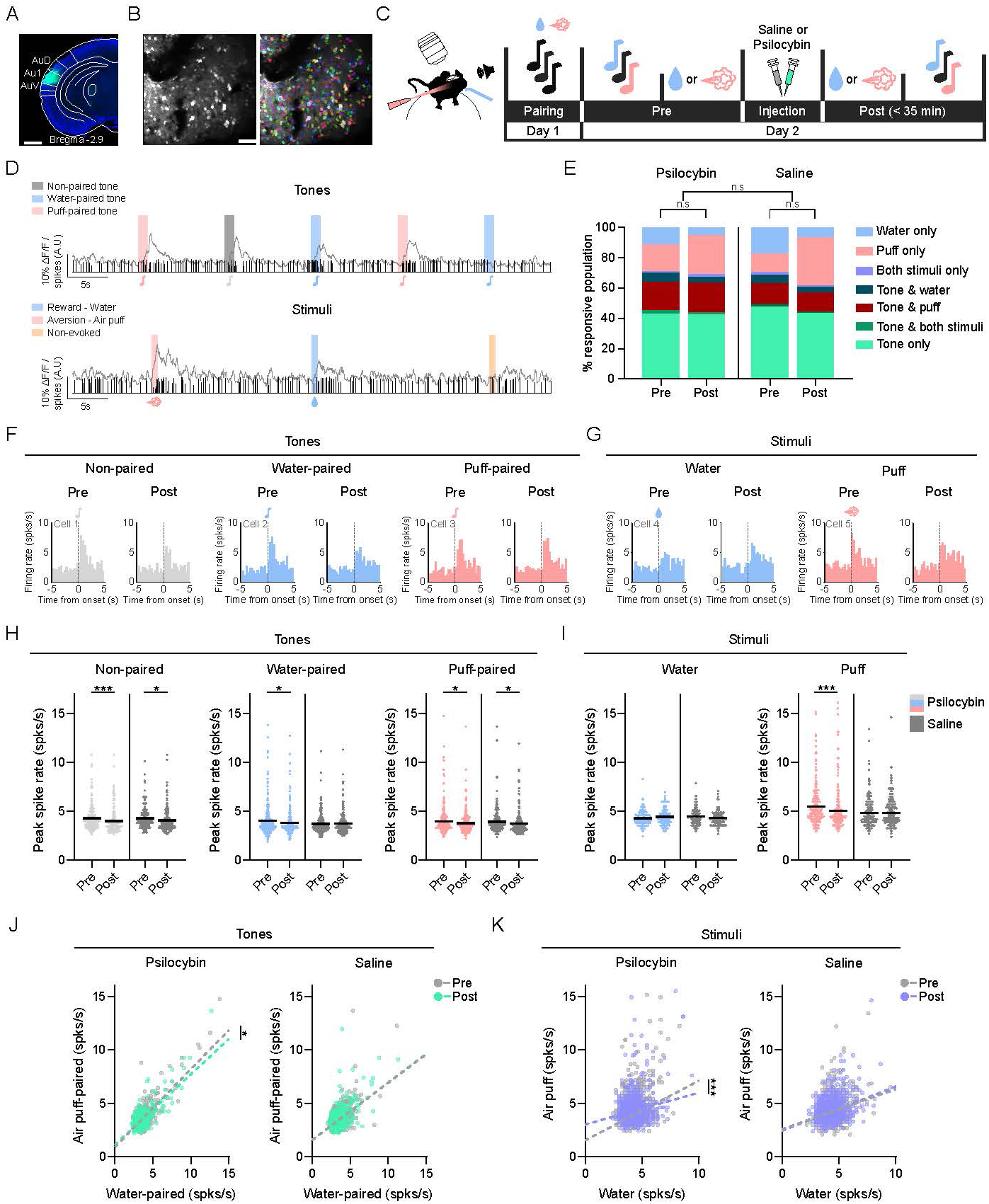
Psilocybin shifts auditory cortex valence responses away from a negative state. Longitudinal in vivo two-photon calcium imaging through a cranial window was used to measure auditory cortex activity in saline and psilocybin (2 mg/kg, i.p.) injected mice. (A) Representative GCaMP8f expression in layer 2/3 of auditory cortex. Scale bar, 1 mm. (B) Example imaging field of view selected by functional mapping. Raw image is shown on the left and cells identified by Suite2P are outlined in the right image. Scale bar, 50 µm. (C) Experimental timeline. (D) Example raw calcium (trace on top) and deconvolved spike (vertical lines) data from a single cell in a mouse when presented with tones (top) and stimuli (bottom). Water/water-paired = blue, air puff/air puff-paired = pink, non-paired = grey, non-evoked = gold. (E) Distribution of neuronal response categories across sessions. Neurons were classified as tone-only, stimulus-only, dual-responsive, or non-responsive based on peri-stimulus spike activity. Compositional data analysis was used to detect shifts in relative response proportions between saline and psilocybin groups across pre and post sessions. n mice: psilocybin: = 5, saline = 5, n cells (pre, post): psilocybin: = 854, 727, saline = 782, 647. (F, G) Example peri-stimulus time histograms for tone (F) and stimuli (G) presentations derived from deconvolved calcium signals in mice given psilocybin. (H) Tone-evoked spike rates grouped by prior valence association, only neurons which had at least a significant increase in tone response across either Pre or Post sessions were included. All data was analyzed using repeated measures mixed models with cell and mouse added as random effects [treatment x sessions x interaction]). Each dot = 1 neuron. Horizontal lines = mean ± SEM. Saline, grey (all), psilocybin, light grey, grey (non-paired), blue (water-paired), pink (air puff-paired). n cells (psilocybin, saline): non-paired 171, 137, water-paired 234, 229, air puff-paired 227, 208. (I) Stimulus-evoked spike rates in response to presentation of water and air puff from neurons which had at least a significant increase in tone response across either Pre or Post sessions. All data was analyzed using a repeated measures mixed model with cell and mouse added as random effects [treatment x sessions x interaction]). Each dot = 1 neuron. Horizontal lines = mean ± SEM. Saline, grey (all), psilocybin, blue (water), pink (air puff). n cells (psilocybin, saline): water 119, 135, air puff 274, 224. (J) Population valence bias for tone-evoked responses. Each point represents one neuron, plotted by its peak spike rate to the water-paired tone versus the air puff-paired tone (pre, grey; post, green). Linear regression lines were fit per group and session, and differences in fitted relationships were assessed using linear regression. Each dot = 1 neuron. Pre = grey, Post = green. n cells (psilocybin, saline): 1189, 1261. (K) Population valence bias for stimulus-evoked responses. Each point represents one neuron, plotted by its peak spike rate responses to water and air puff (pre, grey, post, purple). Linear regression lines were fit per group and session, and differences in fitted relationships were assessed using linear regression. Each dot = 1 neuron. Pre = grey, Post = purple. n cells (psilocybin, saline): 1189, 1261.

### Aversive-only longitudinal dataset paradigm (Figures 2-4)

In Week 1, mice were habituated to head fixation and stimulus delivery, then received an intraperitoneal saline injection followed by pairing of Tone 1 (9 kHz) with air puffs (30-40 pairings; 1 s tone co-terminating with a 0.5 s air puff; 11-17 s pseudorandom inter-trial interval). In Weeks 2-3, mice underwent longitudinal imaging sessions using the same overall block structure as the reward/aversive dataset, but with an updated tone block (9,13 and 22 kHz; 1 s) and an air puff-only stimulus block. A Pre session was acquired the day before injection to establish baseline tone and puff responses. On injection day, mice received intraperitoneal saline or psilocybin. Within 2 minutes of injection, the field of view was re-identified and re-aligned to the single-cell level. An 8-minute non-evoked recording was acquired both before and after re-alignment to capture immediate post-injection activity. The Post session then consisted of a dedicated tone block followed by a dedicated air puff block acquired within 10-35 minutes after injection. Immediately after Post imaging, Tone 3 (22 kHz) was paired with air puffs using the same co-termination structure as Tone 1. Tone 2 (13 kHz) was presented as an unpaired tone throughout the experiment. A Post +1 session was acquired the following day using the same tone and air puff blocks. In Week 3, imaging was repeated at Post +8 (eight days after injection day) to assess persistence.

**Figure 2.**
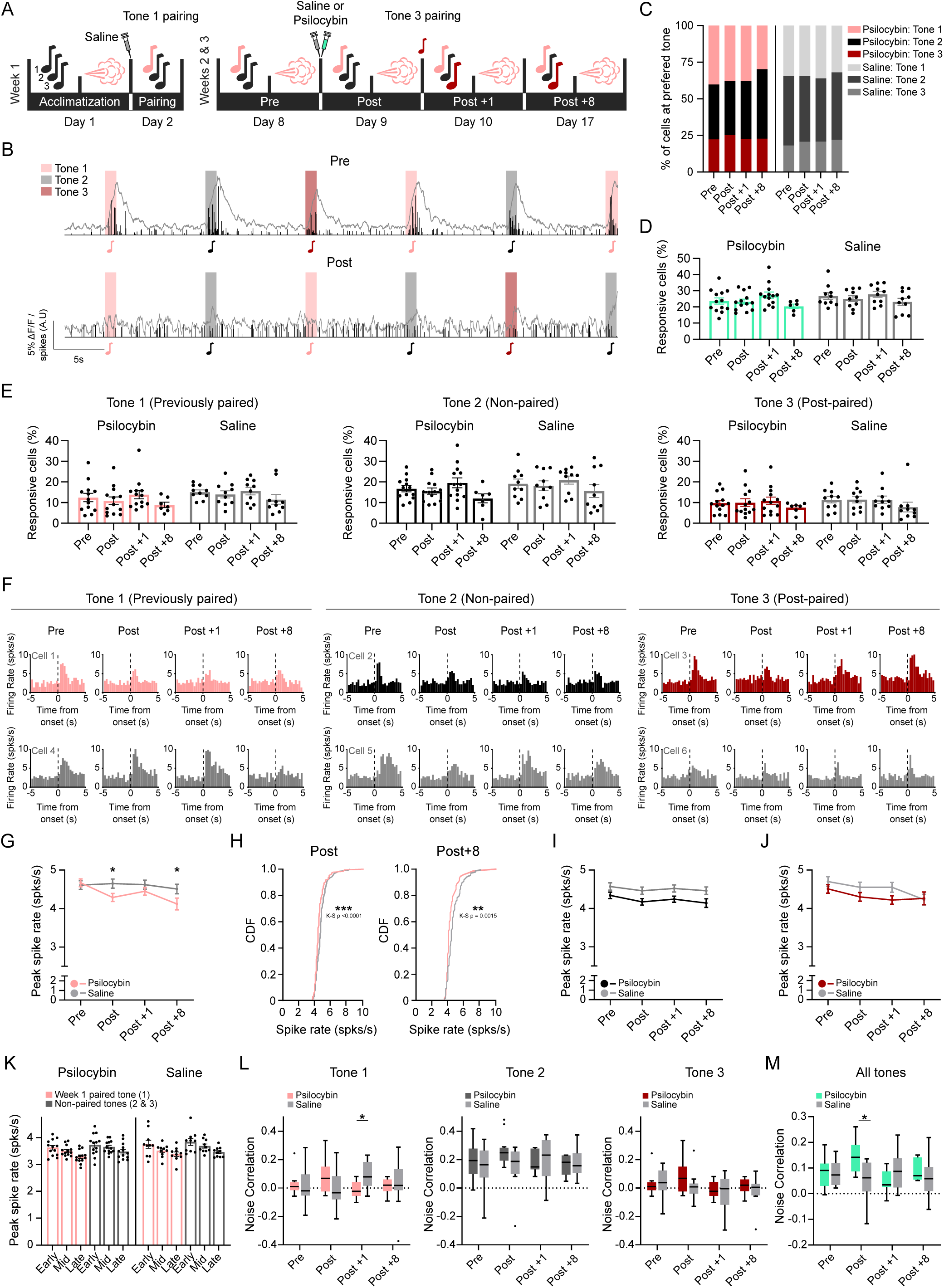
Psilocybin selectively changes responses to previously established aversive associations. Tones were presented in dedicated tone blocks across multiple imaging days. Responses were quantified as peak spike rates from deconvolved spiking as peri-stimulus time histograms (PSTHs) across sessions (Pre, Post, Post +1, Post +8), with tone identity tracked as previously paired (Tone 1), unpaired (Tone 2), or paired during the Post session (Tone 3). (A) Experimental timeline. (B) Example raw calcium and deconvolved spike data from a single cell in a mouse prior (top) to and following (bottom) psilocybin when presented with 1-s tones. (C) Tone preference (based on peak spike rates) distributions across sessions. Compositional analysis was performed using isometric log-ratio transformation with linear mixed-effects modeling of ILR coordinates. (D) All tone population response rates across sessions. Repeated measures mixed-effects models. Each dot represents one mouse. Bars = mean ± SEM. Psilocybin = green, saline = grey. n cells (psilocybin, saline): 2450, 1699. (E) Tone specific population response rates across sessions. Repeated measures mixed-effects models. Each dot represents one mouse. Bars = mean ± SEM. Psilocybin = pink, black, red; saline = grey. (F) Representative PSTH examples of longitudinal tone-evoked spike responses from individual neurons. Top: psilocybin administered, bottom: saline administered. Psilocybin = pink, black, red; saline = grey. (G, I, J) Tone 1 (Previously-paired), Tone 2 (Non-paired) and Tone 3 (Post-paired) peak spike rate responses during weeks 2 and 3. Repeated measures mixed-effects models with post-hoc Holm-Bonferroni comparisons. Lines = mean ± SEM. n cells for all tone specific data (Pre, Post, Post +1, Post +8): psilocybin; tone 1: 294, 240, 286, 81, tone 2: 382, 334, 464, 129, tone 3: 236, 221, 234, 70. saline; tone 1: 206, 188, 208, 160, tone 2: 250, 270, 297, 240, tone 3: 159, 143, 152, 120. (H) Cumulative frequency distributions in response spike rate distributions during the Previously-paired tone Post and Post +8 sessions were assessed using Kolmogorov-Smirnov tests. n cells (Post, Post +8): psilocybin; 240, 188, saline; 81, 160. (K) Trial-stage analysis. Post-session tone responses were divided into early, middle, and late tertiles (approx. 4 minutes each). Repeated measures mixed-effects models tested effects of treatment, trial stage, and interaction. Each dot represents one mouse. Bars = mean ± SEM. n cells (psilocybin, saline): tone 1: 240, 188, tone 2 & 3: 555, 413. (L) Tone-specific noise correlation. Pairwise Pearson correlations were calculated from trial-by-trial response residuals for each tone, and session-level median off-diagonal correlation coefficients were extracted. Repeated measures mixed-effects models. Line = median, whiskers = Tukey’s. n cells (psilocybin, saline): 2161, 1699. (M) Noise correlation across all tones. Pearson correlations were calculated from trial-by-trial response residuals pooled across tones, and session-level median off-diagonal correlation coefficients were analyzed. Repeated measures mixed-effects models. Line = median, whiskers = Tukey’s. n cells (psilocybin, saline): 2161, 1699.

**Figure 3.**
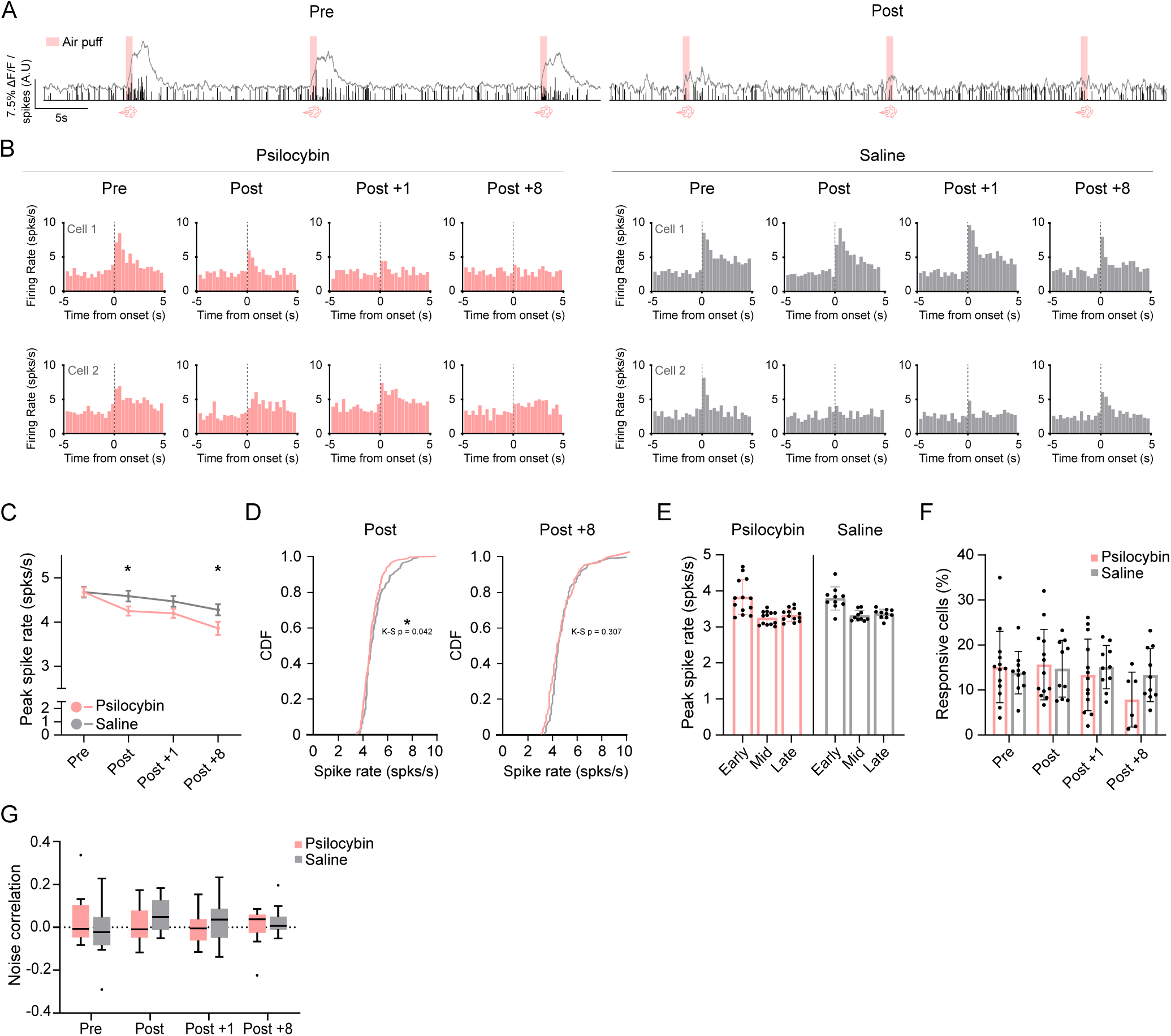
Psilocybin attenuates aversive stimulus-evoked spiking. Air puffs were delivered in a dedicated stimulus block. Responses were quantified as peak spike rates from deconvolved spiking as PSTHs across sessions (Pre, Post, Post +1, Post +8). (A) Example raw calcium and deconvolved spike data of air puff presentations from a single cell in a mouse prior (left) to and following (right) psilocybin when presented with 1-s tones. (B) Example PSTHs aligned to air-puff onset (0.5 s, 15 PSI, ipsilateral eye). (C) Puff-evoked peak spike rates across sessions. Longitudinal effects of treatment, session, and treatment × session were tested using Repeated measures mixed-effects models with random intercepts for mouse and cell. Post-hoc contrasts were corrected for multiple comparisons (Holm-Bonferroni). Lines = mean ± SEM. n = 13 psilocybin, n = 10 saline mice. n cells (Pre, Post, Post +1, Post +8): psilocybin; 414, 446, 368, 86, saline: 236, 250, 262, 229. (D) Cumulative frequency distributions of puff-evoked peak spike rates during the acute Post session. Between-group distribution differences were assessed using two-sample Kolmogorov-Smirnov tests. n cells (Post, Post +8): psilocybin; 446, 86, saline: 250, 229. (E) Trial-order analysis of puff responses within session (early, middle, late trial tertiles). Effects of treatment, trial order, and interaction were tested using Repeated measures mixed-effects models. Each dot = one mouse. Bars = mean ± SEM. n cells (Post): psilocybin; 446, saline: 250. (F) Puff response probability across sessions (fraction of cells classified as responsive). Treatment, session, and treatment × session effects were evaluated using Repeated measures mixed-effects models. Each dot = one mouse. Bars = mean ± SEM. n cells (Pre, Post, Post +1, Post +8): psilocybin; 414, 446, 368, 86, saline: 236, 250, 262, 229. (G) Noise correlation during puff responses. Pairwise Pearson correlations were computed from trial-by-trial residual responses, and session-level median off-diagonal correlation coefficients were analyzed using Repeated measures mixed-effects models. Line = median, whiskers = Tukey’s. n cells (psilocybin, saline): 2450, 1699.

### Calcium Data Processing and Analysis

Data from all imaging sessions for each mouse were processed together as a single dataset in Suite2P to maintain a consistent reference frame and stable ROI identities across days. Suite2P performed motion correction and registration, ROI detection, and fluorescence extraction. Neuropil correction was applied to obtain neuropil-corrected ROI traces, and deconvolution was performed to estimate estimated spikes from the calcium signals [43]. ROIs were curated to exclude non-somatic processes and artefactual signals, and the final ROI set and cell identities were then carried forward for all subsequent per-session analyses using custom MATLAB code. Tone and stimulus timestamps were aligned to the imaging data and used to extract peri-event windows. Peri-stimulus time histograms (PSTHs) were computed from Suite2p deconvolved spike trains using 0.33 s bins. Spike rate (events/s) was computed as the fraction of frames containing spike events (spikes > 0), multiplied by the frame rate. For each cell and condition, trial-wise mean spike rates were computed in pre- and post-event windows and compared using a paired sign-flip permutation test (5,000 permutations; two-tailed, α = 0.05) on the mean paired difference (post - pre); cells with fewer than two valid trials were excluded. Significant positive changes were classified as responsive and significant negative changes as suppressed. Peak responses were defined as the maximum post-event PSTH bin (‘PeakSpike’) and, when applicable, the peak ΔF/F within 3 s of tone or stimulus onset. For the multi-valence cohort, cells were classified as active if they showed a significant response in either the Pre or Post session. For the air puff only cohort, activity was defined on a per-session basis, with cells classified as active only for sessions in which they were significantly responsive. For tone-preference analyses, only neurons that were active in at least two sessions were included.

### Behavioural Tracking and Video Analysis

Mouse movement was recorded using a PhenoSys JetBall laser sensor and quantified as frame-to-frame Euclidean distance. Facial behavior was recorded at 30-Hz using an infrared camera (Basler, acA1300-60gmNIR, with a 12 mm lens) with 850 nm LED illumination (Thorlabs, LIU850A) focused on a side profile ipsilateral to the cranial window and air puff illuminated. Videos were processed using DeepLabCut [44], a Resnet-50 model was trained on approximately 350 manually labeled frames. Eighteen landmarks were placed around the eye, nose, whisker, mouth, and ear points. Blink amplitude was defined as the Euclidean distance between upper and lower eyelids, and facial motion was quantified as frame-to-frame displacement of facial landmarks (nose, whisker, mouth). Behavioral data were temporally aligned to calcium recordings and stimulus onset to generate matching windows to each imaging session.

### Histology and verification of viral expression

At the conclusion of experiments, brains were collected and processed for fluorescence histology to confirm GCaMP expression and targeting. Tissue was fixed, cryo-sectioned coronally, mounted with DAPI-containing medium (Southern Biotech, #0100-20), and imaged by fluorescence microscopy. Expression location was evaluated relative to auditory cortex landmarks/atlas, and off-target cases were excluded (Supplementary Figure 2).

### Statistical Analysis

All data were presented as individual data points and/or expressed as mean ± SEM. Longitudinal neural data were analyzed using linear mixed-effects models implemented in RStudio (2026.01.1). Models primarily used cell-level data and accounted for repeated measurements of neurons nested within mice using random intercepts for ‘Mouse’ and ‘Unique cell’. This design also ensured any drug induced heterogenous effects or extreme variation could be accounted for across cell and mouse levels. Mouse level data is found in Supplementary Figure 3B-D & S4.

The primary models were:

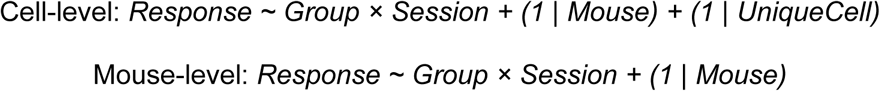

Treatment group and Session (as a number, or factor; to allow for non-linear effects over days to a week) were included as fixed effects. Type III ANOVA with Satterthwaite approximation was used to assess fixed effects, and post-hoc comparisons were corrected using Holm-Bonferroni or Tukey’s method. Significance was set at p = 0.05. Repeated measures mixed models for mouse level data were also performed using RStudio or processed in Graphpad Prism (8 or 10) using a similar mixed model function.

Cumulative distribution functions (CDFs). CDFs were calculated from the raw values for each group/session and plotted to visualize differences in the distribution of spike rates. Kolmogorov-Smirnov tests were performed in MATLAB (kstest2). The maximum distance between CDFs was reported as the K-S statistic (D). Significance was assessed using two-sided tests with p = 0.05.

Compositional data analysis. Because response-category percentages summed to 100% and are therefore not independent, we analyzed these data using compositional data analysis (CoDa) [45]. Each analysis was run separately for each response-category set and, where applicable, by modality. For each mouse and session, category percentages were treated as a D-part composition and transformed using an isometric log-ratio (ILR) transform (R package compositions), which maps the composition into D-1 orthonormal coordinates (V₁ to V_{D-1}) in Euclidean space suitable for standard mixed-effects modeling. Each coordinate V_i represents a log-ratio balance contrasting subsets of categories, such that changes reflect redistribution across categories rather than shifts in any single one. The same ILR basis was applied across all mice and sessions within a given analysis, making each V_i directly comparable across groups and sessions. We fit linear mixed-effects models to each ILR coordinate using:

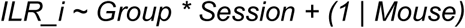

This treated Group and Session (factor) and their interaction as fixed effects and including a random intercept for Mouse. Fixed effects were evaluated with Type III Wald χ² tests, and when appropriate, session-wise post hoc contrasts were performed using estimated marginal means with Bonferroni correction. For interpretability, we back-transformed model predictions from ILR space to proportions and computed correlations between each V_i and the original category proportions to identify which response classes most strongly contributed to each balance. Group-level differences in the full composition were additionally summarized using Aitchison distance (Euclidean distance in ILR space) with 10,000 iteration permutation testing. All CoDa data were run using RStudio using the ‘compositions’, ‘lme4’, ‘lmerTest’, ‘car’ and ‘emmeans’ packages.

Noise Correlation Analysis. Noise correlations were calculated to measure how similarly pairs of neurons fluctuated from trial to trial after removing the average stimulus response. For each neuron and trial, we computed a single “trial response” as mean activity in the response window minus mean activity in the baseline window. In tone sessions, trials were grouped by frequency, and we removed each neuron’s average response for that frequency (demeaning within frequency) to get residuals that reflect trial-to-trial variability rather than tuning. We then computed Pearson correlations between pairs of neurons using only trials where both neurons had valid values (pairwise-complete), requiring a minimum number of shared trials. For each session, the noise correlation was summarized as the median of the off-diagonal pairwise correlation values. Full statistics and resources can be found in the Statistics Table and Resources Table.

## Results

### Psilocybin shifts auditory cortex valence responses away from a negative state

To determine whether tone responses and valence-related information are modified in the auditory cortex by psilocybin administration, C57BL/6 mice were injected with Syn-AAV-GCaMP8f and implanted with cranial windows over auditory cortex (Figure 1A-B). Across all imaging sessions mice were awake and head fixed. Five weeks after window surgeries, tonotopic mapping was carried out to locate the auditory cortex along with acclimatization to the imaging stage. Mice then underwent water restriction before the pairing of tones with valenced stimuli: water (rewarding) and air puff (aversive) (Figure 1C). 8-kHz and 13-kHz tone were randomly assigned to either water or air puff in each mouse. The following day, neurons within the auditory cortex were repeatedly imaged before (Pre) and after (Post) intraperitoneal injection of either psilocybin (2 mg/kg in 0.1 mL saline; n = 5 mice, 2 females, 3 males) or saline alone (0.1 mL; n = 5 mice, 3 females, 2 males). No sex-dependent differences were detected for tone or stimulus responses (Supplementary Figure 3A), and male and female mice were merged into a single cohort for data analyses.

During the Pre and Post sessions, the same auditory cortex neurons were imaged during two blocks. Block 1 consisted of interspersed randomly ordered, 1-s pure tones across 3 frequencies (4, 8, 13 kHz). Block 2 consisted of pseudorandom interspersed valence stimuli trials (water: 5 µl per trial; air puff: 0.5-s puff duration, 15-PSI). Within Block 2, the stimulus was omitted on every third trial to measure spontaneous activity. All Post session data were acquired within 35 minutes of injection, to correspond with the psilocybin active phase and peak head twitch response (HTR) period in mice [41,43]. Calcium signals from all sessions of each animal were processed as a single dataset using Suite2P to ensure consistent ROI tracking across sessions and were subsequently deconvolved to infer estimated neural spiking (Figure 1D). The resulting spike data were used to construct peri-stimulus time histograms (PSTHs) aligned to each tone or stimulus. We assessed the effect of psilocybin on evoked spiking by extracting the peak spiking rate from PSTHs within a 3 s window following tone or stimulus onset compared to the corresponding pre-onset 3-s baseline within each session.

Across both saline- and psilocybin-treated mice, neurons exhibited tone-only, stimuli-only, and combined tone-stimuli responses during both Pre and Post sessions. The plurality of neurons responded to tones only (Figure 1E; Pre: PSY = 43.1 ± 3.42%, SAL = 47.7 ± 4.17%; Post: PSY = 42.7 ± 8.51%, SAL = 43.4 ± 4.98%). The remaining tone-responsive neurons responded to both tone and stimuli, with tone-and-puff responsive neurons representing the largest population within this category (Figure 1E; tone & air puff; Pre: PSY = 18.6 ± 3.38%; SAL = 13.5 ± 4.03%; Post: PSY = 20.0 ± 4.62%, SAL = 12.7 ± 1.17%). Among stimulus-only neurons, a larger proportion responded to air puff than to water or to both stimuli across both treatment groups and sessions, except the saline Pre session (Figure 1E; air puff only; Pre: PSY = 17.6 ± 1.75%; SAL = 12.8 ± 3.52%; Post: PSY = 25.79 ± 5.33%, SAL = 31.45 ± 4.16%). However, using compositional data analysis, no proportional differences were recorded between categories across treatments and sessions, indicating psilocybin does not shift the distribution of response categories (Figure 1E).

For tone-evoked spiking, both saline and psilocybin mice had reduced Post spike rates to non-paired tones and tones paired with air puffs, (Figure 1F, 1H; non-paired: session p < 0.0001, water-paired: all n.s., puff-paired: session p < 0.0001). The similar reductions in spike rates following psilocybin and saline injections suggest that the injection procedure itself likely caused a general decrease in tone-evoked responses. For stimulus-evoked spiking, psilocybin, but not saline, had significantly reduced Post puff-evoked spike rates (Figure 1G, 1I; air puff: session p = 0.0011, treatment x session p = 0.00103). Neither saline nor psilocybin significantly shifted spike rates to water stimulus (Figure 1G, 1I). Within the stimulus block, non-evoked activity did not differ across treatment or session (Supplementary Figure 3D). Together, these findings indicate that psilocybin selectively reduced puff-evoked spiking.

To assess whether psilocybin alters the population level balance between positive and negative valence responses, we fit linear regressions relating water and puff evoked spiking across all neurons for each treatment and session. Psilocybin injection shifted the population spike rate away from puff paired tones (Figures 1J; psilocybin: p < 0.0001) and puff (Figure 1K; psilocybin: p = 0.0008), whereas no such change was measured in saline injected mice. These findings indicate that psilocybin reorganizes population level valence encoding, biasing spike rates away from puff and puff paired tones.

Together, these results indicate that psilocybin alters auditory cortex responses to valenced stimuli, reducing puff-evoked spike rates at the single-cell level while biasing population spiking away from puff and puff-paired tones.

### Psilocybin selectively changes responses to previously established aversive associations

Our results indicate that psilocybin selectively reduces spiking responses to aversive stimuli. However, because the injection procedure produced a general decrease in tone responses, we were unable to assess whether psilocybin also influences responses to tones previously paired with aversive stimuli. To address this question, we modified the study design to perform post-injection imaging on the same day as well as 1 and 8 days after administration (Figure 2A). In addition, tones were paired with air puffs one week prior to psilocybin treatment, and tones and air puffs were presented in separate blocks during the imaging session. This approach allowed imaging after the acute effects of the injection had worn off, permitted longer consolidation of the tone-puff association, enabled examination of how responses to aversive stimuli evolve over time, and avoided potential interactions between tone- and puff-evoked responses. In this design, tone 1 was paired with air puff following saline injection during week 1. During week 2-3, mice underwent Pre, Post, Post+1 (day) and Post +8 imaging. Before the Post session, approximately half of the mice received psilocybin (2 mg/kg; n = 13 mice, 7 females, 6 males) and the remainder received saline (n = 10 mice, 6 females, 4 males). Following the Post session imaging, tone 3 was paired with air puffs, while tone 2 remained unpaired throughout. Imaging was repeated one week later to assess longer-term effects in the Post +8 session. No sex-dependent differences were identified (Supplementary Figure 5), so data from all male and female mice were pooled for analysis. Mice showed robust tone responsiveness across the 3 tones presented (Figure 2B). To determine whether tuning preference was shifted by psilocybin, tone preference distributions were calculated across all sessions. No significant treatment or sessional effects were observed (Figure 2C: Compositional Data Analysis, χ², all p > 0.05). The proportion of tone responsive cells were also stable across groups for all tones (Figure 2D & 2E).

We next examined the tone-specific spike rates across treatment and sessions. Spiking evoked by tone 1, which had been paired with air puff one week prior to psilocybin administration, showed a progressive decrease following psilocybin treatment over time in both the saline and psilocybin groups, with a greater decreased in the psilocybin group (Figure 2F, 2G; session p = 0.0034, treatment x session p = 0.0121, Post-hoc saline vs. psilocybin: Post p = 0.0297, Post +8 p = 0.0478). The cumulative spike rate distribution in the Post and Post+8 sessions shifted toward lower rates in the psilocybin group compared with the saline group (Figure 2H; saline vs. psilocybin CDF, Kolmogorov-Smirnov test, Post: p < 0.0001, Post +8: p = 0.0015). Spiking evoked by unpaired tone 2 and by tone 3, which was paired during the Post session within 35 minutes of injection, did not differ between the psilocybin and saline groups across all sessions (Figure 2I & 2J).

A recent study has showed that psilocybin produces biphasic changes in tone-evoked calcium transients in the auditory cortex, i.e. an early increase starting 5 minutes of the injection then a decrease starting 30 minutes later [41]. We examined whether a similar biphasic effect was also present in our dataset by examining responses to tones during the Post session prior to tone 3 pairing with air puffs in week 2. In our paradigm tone presentation started 10 minutes after injection, however, prior to this, non-evoked spike rates were measured but unchanged by psilocybin (Supplementary Figure 6A). We then split the tone trials in the Post session into equal tertiles around 4 minutes each, representing early, middle and late trials. Spike rates declined across early, middle, and late-stage trials for both previously paired and unpaired tones, with no differences between the saline and psilocybin groups (Figure 2K; paired-tone: trial stage p < 0.0001, trial stage x group interaction p = 0.331, unpaired-tones: trial stage p < 0.0001, trial stage x group interaction p = 0.7509).

We then assessed population-level coordination using noise correlation analysis. Noise correlations quantify trial-to-trial co-variability in neurons’ responses after removing the mean stimulus-evoked component. Higher noise correlation indicates greater shared variability across neurons, consistent with stronger common input and or tighter functional coupling within the population. When tones were analyzed separately, a treatment-by-session interaction was detected only for tone 1, the previously puff-paired tone, driven by reduced noise correlation in psilocybin-treated mice at Post+1 (Figure 2L; Tone 1 treatment × session p = 0.0265, post-hoc saline vs. psilocybin: Post+1 p = 0.0393). These results suggest that psilocybin attenuates coordination among cells responding to tones previously paired with aversive stimuli. Across all tone responsive cells, noise correlation also showed a treatment-by-session interaction, with psilocybin mice exhibiting higher noise correlation during the acute Post session (Figure 2M; all tones, treatment × session p = 0.0019, post-hoc saline vs. psilocybin: Post p = 0.0381). Taken together, these findings suggest that psilocybin induces an acute, network-wide increase in coordination across tone-responsive cells, while later reducing coordination specifically within neurons responding to negatively paired tones. This pattern is consistent with a transient shift toward more diffuse synchrony during the acute phase, followed by selective decoupling of aversive-associated representations. Together, these findings indicate that psilocybin specifically modifies spiking responses to tones that were paired with aversive stimuli before treatment, but not to tones paired afterward or unpaired tones.

### Psilocybin attenuates aversive stimulus-evoked spiking

Auditory cortex neurons exhibited robust air-puff–evoked responses in naïve animals and across subsequent sessions in both saline- and psilocybin-treated mice (Figure 3A). Puff-evoked spike rates decreased across post-injection sessions in both groups, with a larger reduction in psilocybin-treated mice (Figure 3B, 3C; session p < 0.0001, treatment x session p < 0.01, post-hoc saline vs. psilocybin: Post p = 0.0478, Post +8 p = 0.0371). Cumulative frequency analysis for the Post but not the Post +8 session showed a spike-rate distribution shifted toward lower values (Figure 3D; CDF: saline vs. psilocybin Kolmogorov-Smirnov test, Post: p = 0.042, Post +8: p = 0.306). The evoked spike rates declined across early, middle, and late trials regardless of treatment, with no differences between the psilocybin and saline groups (Figure 3E; trial stage p < 0.0001), indicating that psilocybin has no effect on the temporal dynamics of spiking responses across trials. The proportion of puff-responsive neurons was comparable between psilocybin and saline groups (Figure 3F). Noise correlation in the puff block was unchanged across sessions and treatments (Figure 3G). These results indicate that in addition to attenuating spiking to tones previously associated with aversive stimuli, psilocybin also blunts spiking to the aversive stimulus itself, without altering the proportion of neurons responding to air puff or their neural synchrony at the population-level.

As locomotor movement has been shown to heterogeneously modify auditory cortex activity, we tested whether locomotor changes could be associated with the psilocybin induced neural activity changes [44]. Locomotor activity during head-fixed imaging was assessed from movement on the ball. Total movement within the post-onset 3-s did not differ between treatment groups across sessions for tones or for air puff (Supplementary Figure 7). To monitor facial activity, mice were recorded with an infrared camera during all imaging sessions. The videos were analyzed using DeepLabCut to track blink responses and facial features (Supplementary Figure 8A & B). Blinks were measured as the Euclidean distance between the top and bottom eyelids. Blinks evoked by co-presentation of tone and air puff during pairings across week 1 and week 2 showed comparable amplitude between psilocybin and saline groups (Supplementary Figure 8C). Air puff-and tone-evoked blink amplitude and latency were stable across treatment groups over all imaging sessions (Supplementary Figure 8D-F). Facial motion metrics derived from whisker, nose, and mouth movements were used to assess overall facial activity and as a proxy for arousal and non-auditory sensory activity. Neither the mean facial motion nor the correlation of facial motion with the peak spike rates differed between psilocybin- and saline-treated mice during either tone or air puff presentations (Supplementary Figure 8L-O). Due to head fixation, no head twitch responses were observed in videos through manual assessment. Together this data suggests psilocybin does not induce any major behavioral changes in locomotion, blink response or overall facial activity in our head fixed setup. Movement, therefore, does not explain the changes in spike rates recorded.

### Psilocybin selectively dampens evoked spiking of puff-only neurons

Psilocybin’s effects appear to be heterogenous across neuronal population within the same region [14]. We recorded heterogeneous neural responses to tones and air puffs, including tone-only, stimulus-only, and dual-responsive neurons that responded to both tones and air puffs (Figure 4A). We then tested whether psilocybin has different effects on these subpopulations. Across sessions, approximately 50 to 70 percent of responsive neurons were tone-only, 15 to 25 percent were dual-responsive, and 10 to 25 percent were stimulus-only. Through compositional analysis, a redistribution was detected in psilocybin-treated mice over time, characterized by a relative increase in tone-only neurons, with the strongest divergence at Post +8, while the puff-only and dual-responsive populations remained comparatively stable with respect to each other (Figure 4B, Compositional Data Analysis: treatment × session interaction: χ² = 10.38, p = 0.016, Tone-only vs others: Post-hoc saline vs. psilocybin: Post +8 p = 0.0309). Saline mice remained stable across all response categories. During tone presentation blocks, both tone-only and dual-responsive neurons showed no differences across treatment and all imaging days (Figure 4C & 4D). For stimulus-evoked responses, air puff-only neurons showed a significant psilocybin induced spike rate decrease over time (Figure 4E, Session p < 0.001, interaction p < 0.01), resulting in a significant reduction at Post +8 in psilocybin-treated animals (Post hoc psilocybin vs. saline, p = 0.0161). Dual-responsive showed no changes across treatment (Figure 4F). Taken together, these analyses suggest that psilocybin-related changes are not uniformly distributed across auditory cortical neurons. Psilocybin induced attenuation appears most evident in aversive stimuli responsive only neurons.

**Figure 4.**
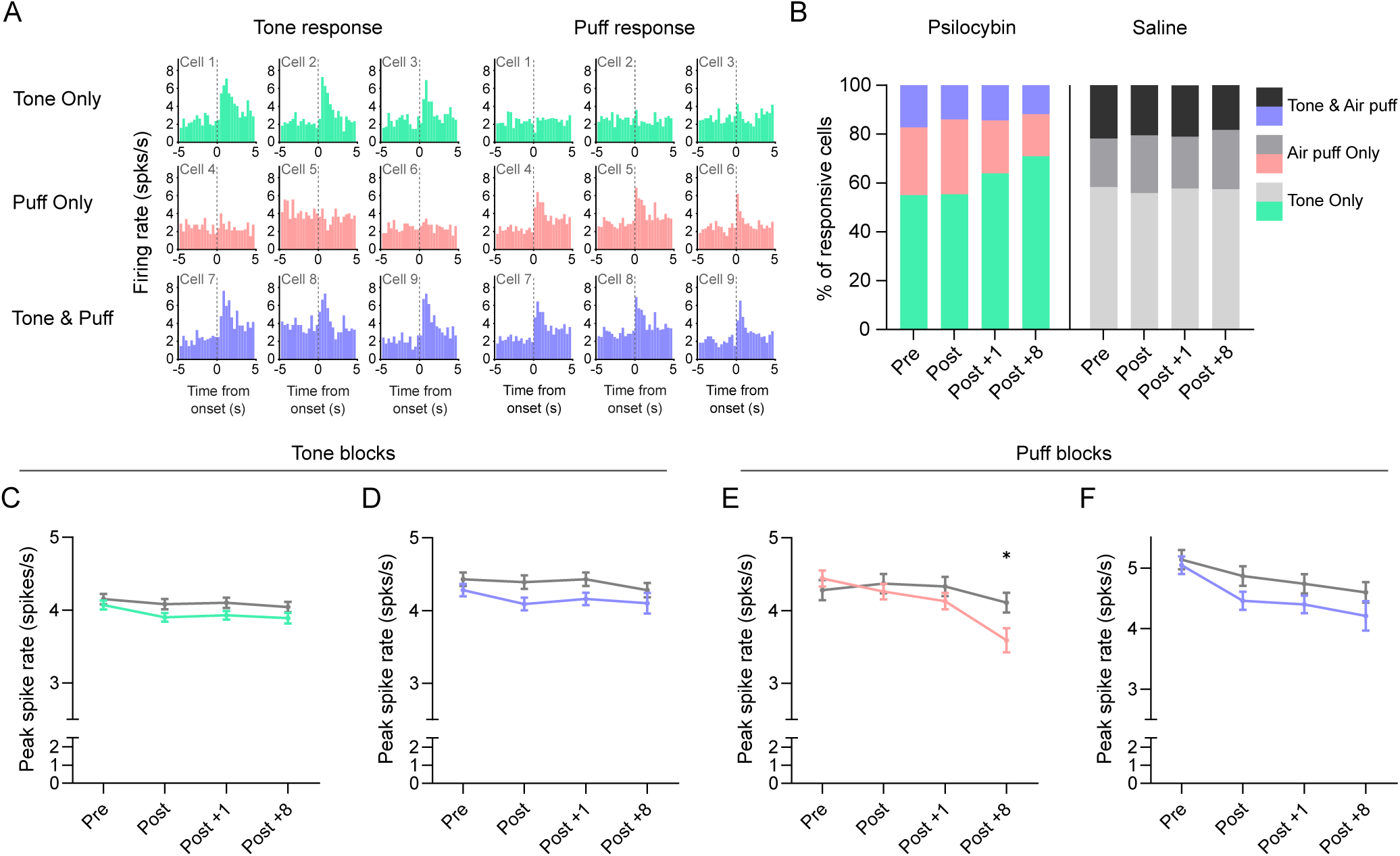
Psilocybin selectively dampens evoked spiking of puff-only neurons. Neurons were stratified by functional responses to identify subpopulations that could contribute disproportionately to psilocybin-related changes in auditory cortex activity. Cells were classified as tone-only (any tone), air puff-only, or dual-responsive (responding to both tones and air-puffs) using peak spike rates from deconvolved spiking-derived PSTHs across sessions (Pre, Post, Post +1, Post +8). (A) Representative PSTHs from example neurons classified as tone-only, stimulus-only, or dual-responsive. (B) Population distribution of response classes across sessions in saline-and psilocybin-treated mice. Compositional analysis was performed using isometric log-ratio transformation with linear mixed-effects modeling of ILR coordinates. n = 13 psilocybin, n = 10 saline mice. n cells (Pre, Post, Post +1, Post +8 totals): psilocybin; 930, 951, 968, 291, saline; 567, 560, 594, 524. (C–D) Tone-evoked responses. Puff-evoked peak spike rates from tone-only (C) and dual-responsive (D) neurons across sessions. Longitudinal effects of treatment, session, and treatment × session were tested using Repeated measures mixed-effects models with random intercepts for mouse and cell. Post-hoc contrasts were corrected for multiple comparisons (Holm-Bonferroni). Line = mean ± SEM. n cells (Pre, Post, Post +1, Post +8): tone-only; psilocybin, 516, 505, 600, 205, saline, 331, 310, 332, 295. Tone and air puff; psilocybin, 175, 135, 164, 31, saline, 128, 123, 135, 105. (E–F) Stimulus-evoked responses. Puff-evoked peak spike rates from air puff-only (E) and dual-responsive (F) neurons across sessions. Longitudinal effects of treatment, session, and treatment × session were tested using Repeated measures mixed-effects models with random intercepts for mouse and cell. Post-hoc contrasts were corrected for multiple comparisons (Holm-Bonferroni). Line = mean ± SEM. n cells (Pre, Post, Post +1, Post +8): air puff-only; psilocybin, 239, 311, 204, 55, saline, 108, 127, 127, 124. Tone and air puff; psilocybin, 175, 135, 164, 31, saline, 128, 123, 135, 105.

## Discussion

Growing evidence suggests that psilocybin may alleviate chronic and often treatment resilient psychiatric conditions marked by persistent negative affect and aberrant emotional learning, including depression, anxiety, and PTSD [6,9]. However, how psilocybin influences neural responses to negatively valenced sensory stimuli remain unclear.

Emotionally salient activity in the auditory cortex has been reported in both humans and rodents [29-31,33,48]. However, relatively little work has directly tested whether auditory cortical activity encodes valence or other extra-auditory variables at the single-cell or ensemble level. In this study, we investigated this question by combining tone and valenced stimulus presentations with longitudinal two-photon imaging of auditory cortex neurons before and after psilocybin administration. By re-imaging the same neurons across weeks, we could attribute changes to treatment and time rather than to differences in which cells were recorded. Among the cells excited by tone and stimuli, roughly 70-80% of them were tone responsive, and about 30-40% responded to stimuli (water and/or air puff), with a distinct subpopulation showing dual responsiveness to both tones and aversive stimuli. Very few cells (< 3%) showed responses to both water and air puffs, with or without tone responsiveness. Across the presented frequencies, both the proportion of tone-responsive neurons and the magnitude of tone-evoked responses were within the range reported in the literature, accounting for methodological differences across studies [49-51]. These findings indicate that individual auditory cortical neurons show valence-associated activity, and that this activity is consistent with growing evidence that auditory cortex represents more than acoustic information alone, incorporating choice-related signals, reward-related variables, and fear-related activity [34,52-54].

Psilocybin did not change the overall distribution of response categories; however, it reduced the evoked spikes rates to air puff and tones paired with puff, but not positively valenced stimulus (water). The effect on tones was specific to the tone that had been paired with the aversive stimulus one week before treatment, but not tones paired 24 hours before, never paired, or paired within 35 min of psilocybin treatment. These changes contribute to an overall population response shift away from aversive tones or stimuli. These findings suggest that although psilocybin can induce rapid spine growth and other plasticity-related changes within hours of administration [3,14,18], it selectively decouples existing, consolidated tone and valence association but not the tone and valence pairing process. The selective effect of psilocybin on consolidated aversive associations is consistent with evidence that psilocybin may be particularly relevant for chronic disorders such as PTSD and major depression, in which persistent maladaptive emotional memories are resistant to change. This interpretation also helps explain why psilocybin’s effect on shorter or early post-conditioning paradigms often produce mixed results: in a multi-institutional study psilocybin reliably reduced acute fear expression but did not yield a reproducible enhancement of extinction 24-48 hours later [20].

The parallel attenuation of evoked spike rates to both puff and the previously puff-paired tone is consistent with a mechanism that shared aversive drive into auditory cortex being weakened by psilocybin. Psilocybin can produce this effect through multiple pathways. The auditory cortex receives aversive, reward, and contextual information from corticostriatal pathways and projections from the basolateral amygdala, anterior cingulate cortex, orbitofrontal cortex, medial geniculate body, and retrosplenial cortex [19,34,48,55-58]. Because psilocybin is known to modulate activity across several limbic and associative nodes (e.g.), the reduced responses to aversive tones and stimuli, but not neutral or rewarding tones, may reflect altered long-range affective input, rather than a uniform change in auditory cortical tone processing. Aversive signals can also modify cortical responsiveness locally. Notably, associative fear learning in the auditory cortex can be implemented through a layer 1 disinhibitory microcircuit that gates CS-US convergence in layer 2/3 [52]. Moreover, given that psilocybin is converted to psilocin and acts primarily through serotonergic signaling, serotonergic input may provide an additional link between psilocybin and the auditory cortex. Consistent with this, serotonergic projections from the raphe nuclei to auditory cortex are known to modulate auditory processing, although the underlying circuit mechanisms and functional consequences remain incompletely understood [59-61].

Our data show that in addition to evoked spike rates at the single-cell level, psilocybin also decreased noise correlation at the population for neurons responsive to tones paired with air puff one week before treatment. This may reflect a change in network state and shared variability that could influence how context is weighted or updated [62]. This possibility is consistent with the relaxed beliefs under psychedelics (REBUS) model, which proposes that psychedelics reduce the influence of strongly weighted priors encoded in higher-order and subcortical circuits, thereby altering how sensory information is integrated [63]. Within this framework, stronger prior associations may be more susceptible to modification, whereas affective states or new associations formed during the acute drug state may not achieve sufficient stability to drive lasting cortical changes. Accordingly, the preferential effect for aversive associations reported here may reflect differences in prior strength or stability rather than valence specificity per se.

Overall, our findings suggest that psilocybin produces selective and time-dependent changes in aversive representations within the auditory cortex. The observed effects appear to preferentially involve previously established negative associations, while sparing basic auditory processing and newly acquired associations. Such specificity may help explain how psilocybin can modify maladaptive emotional representations without broadly disrupting sensory function. Psilocybin induces a broadly heightened, emotionally labile state that may extend well beyond the dosing session. This effect is a significant concern associated with psilocybin-assisted psychotherapy. Our data suggests that transient negative experiences arising within this window are not preferentially strengthened or persistently encoded. This is consistent with two recent studies showing that peritraumatic exposure to psychedelics does not systematically exacerbate aversive memories. Netzer et al. examined PTSD outcomes among survivors of a large-scale mass trauma event experienced under psychoactive substances. Survivors who had consumed psilocybin or LSD at the time of the attack did not show elevated PTSD symptom severity relative to those drug-free [64]. Barnir et al. reported complementary findings in a separate cohort, showing that psychedelic use was associated with significantly lower post traumatic symptoms relative to other substances or no substance use [65]. Collectively, our findings and prior work support the safety and continued development of psilocybin assisted psychotherapy. A previous study reported a biphasic modulation of tone-evoked calcium responses in A1 layer 2/3 after 2 mg/kg psilocybin, with enhanced response amplitude beginning around 5 minutes after dosing followed by reduced amplitude beginning around 30 minutes after dosing [41]. We did not observe treatment-specific biphasic changes. This discrepancy may reflect differences in sampling and readout. Our tone-evoked analyses began 10 minutes post-injection, overlapping part of the reported early phase but not capturing the earliest post-injection period, and they did not extend into the later attenuated phase. In addition, we quantified deconvolved spiking rather than calcium transient amplitude, and our recordings were acquired during an aversive-learning context that can strongly modulate sensory cortical gain. Engagement of threat-related circuits during aversive stimulation and conditioning is known to exert strong top-down modulation of sensory cortical processing [23,52]. Whether psilocybin’s acute effects interact with behavioral state, including attention and arousal, remains to be directly addressed in future.

Several important questions emerge from the present findings that will guide future work. Our recordings were restricted to layer 2/3 auditory cortex neurons without distinction between neuronal subtypes or projection-defined populations, leaving open the question of whether psilocybin’s effects are broadly distributed or concentrated within specific cell classes. Deeper cortical layers, particularly layers 5/6, are more strongly associated with serotonergic signaling and represent the primary source of long-range cortical output, making them compelling targets for understanding how psilocybin’s effects may propagate across circuits. Recent work implicating distinct cortical populations in psychedelic action [14,66] suggests that cell-type resolution will be important for interpreting these effects. Future studies combining longitudinal imaging with genetic labelling or retrograde tracing would clarify whether psilocybin preferentially acts on specific neuronal subtypes, and whether its effects reflect local intracortical dynamics or are driven by long-range inputs from valence-related regions such as the basolateral amygdala or prefrontal cortex. The head-fixed preparation and stimulus parameters employed here prioritized proper experimental control and imaging stability but constrained the range of behavioral states and motivational contexts sampled. Whether the observed valence-specific effects generalize across intensity, modality, or unconstrained behavioral conditions remains to be established. Psilocybin’s influence on positive valence encoding perhaps requires stronger or more naturalistic elements. Future work employing freely moving paradigms, graded reinforcer intensities, or multimodal aversive stimuli will be important for establishing the boundary conditions of the present findings. Finally, all experiments were conducted in C57BL/6 mice aged 18–22 weeks. Although significant hearing loss in this strain emerges later and at higher frequencies than those used here [67], replication in hearing-preserved strains such as CBA/CaJ would further strengthen confidence in the auditory specificity of these findings.

## Supporting information

Statistics Table

Resources Table

## Acknowledgments

Contributions of the NIH authors were made as part of their official duties as NIH federal employees, are in compliance with agency policy requirements, and are considered Works of the United States Government. The findings and conclusions presented in this paper are those of the authors and do not necessarily reflect the views of the NIH or the US Department of Health and Human Services. We also thank George Dold, David Ide, and the Section on Instrumentation of the NIH for fabricating the behavioral apparatus.

## Author contributions

J.J and Z.L conceived the study, designed experiments and wrote the manuscript. J.J and R.T performed cranial window surgeries. J.J performed two-photon imaging experiments, script writing and all analysis. Y.E collected tissue, performed sectioning and imaging.

## Data Availability Statement

Data can be available on request

