## Supplemental data for "Psilocybin Attenuates Cortical Representations of Aversion in the Mouse Auditory Cortex"

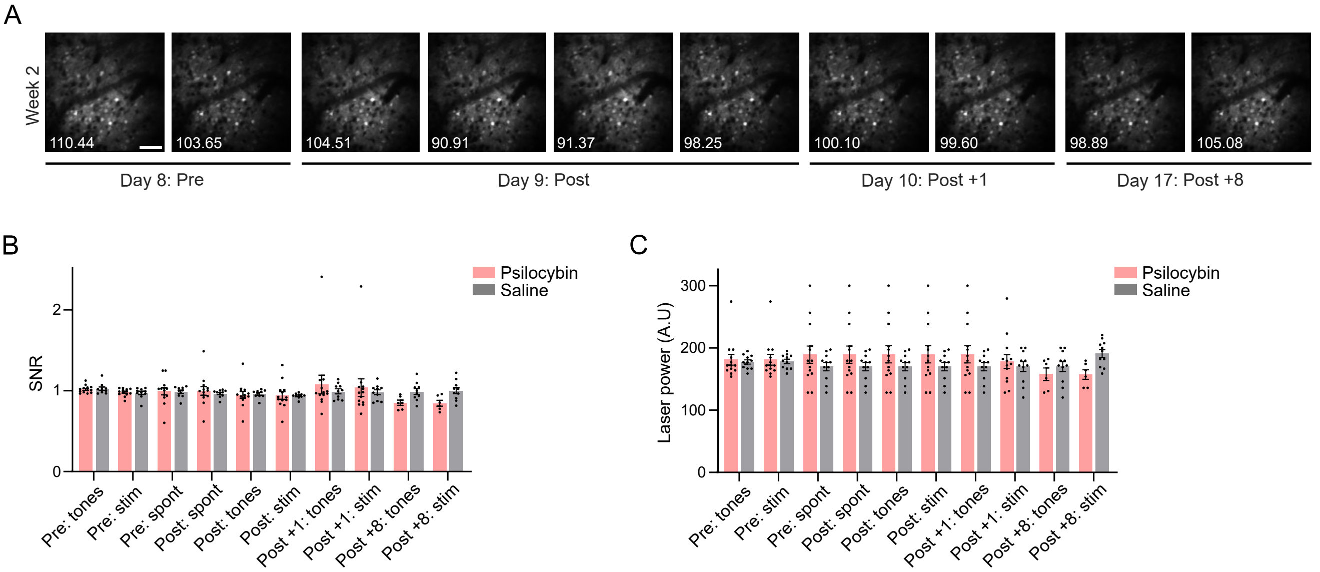


**Supplementary Figure 1. Imaging quality and session consistency**

Session quality was evaluated using mean projection images and percentile-based signal-to-noise ratio (SNR) estimates, with signal defined as pixels above the 95th percentile and noise as pixels below the 5th percentile. SNR was normalized to the average of pre: tones and pre: stim. Each dot = one mouse. Bars = mean ± SEM. Per panel n = 13 psilocybin, n = 10 saline mice. Repeated measures mixed-effects model [treatment x session x interaction], post-hoc Tukey's test for all data. (A) Maximum intensity projections of a representative imaging field of view from a psilocybin-treated mouse across all sessions, with SNR values shown for each session. Scale bar = 50 μm. (B) Mean SNR values across sessions, averaged over all mice. Each dot = 1 mouse. psilocybin = pink, saline = grey. (C) Average laser power (arbitrary units) used per mouse across sessions. Each dot = 1 mouse. psilocybin = pink, saline = grey.

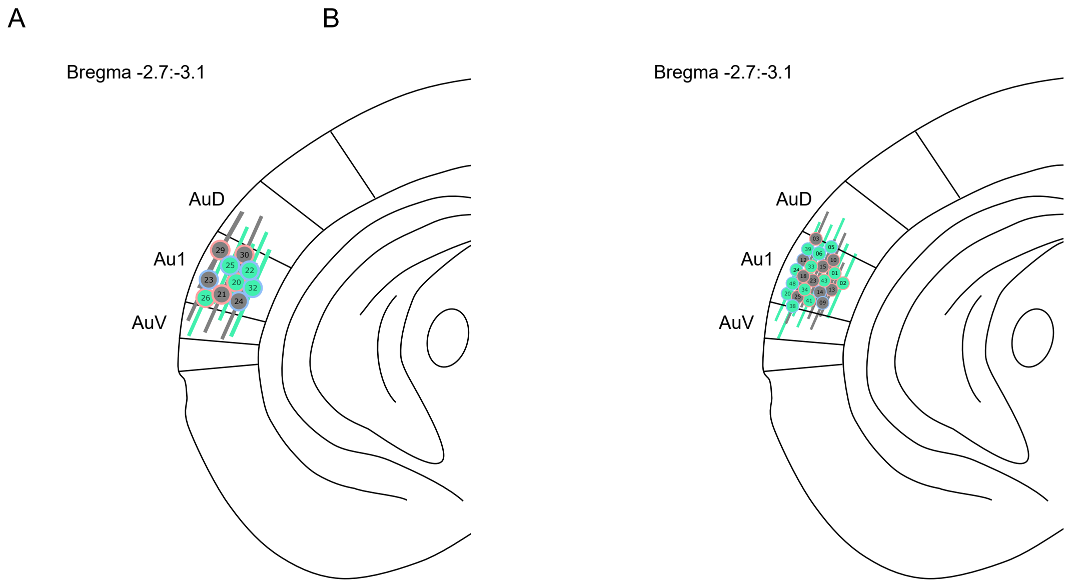


**Supplementary Figure 2. GCaMP expression in all mice**

GCaMP8f expression was qualitatively verified following perfusion and imaged via 30µm post-fixed brain sections. Mice are labeled by internal color indicating treatment group (psilocybin = green; saline = grey) and an external ring indicating sex (male = blue; female = pink). (A) Expression sites per mouse, gender and treatment for mice used in Figure 1. n = 5 psilocybin, n = 5 saline mice. (B) Expression sites per mouse, gender and treatment for mice used in Figures 2-4. n = 13 psilocybin, n = 10 saline mice.

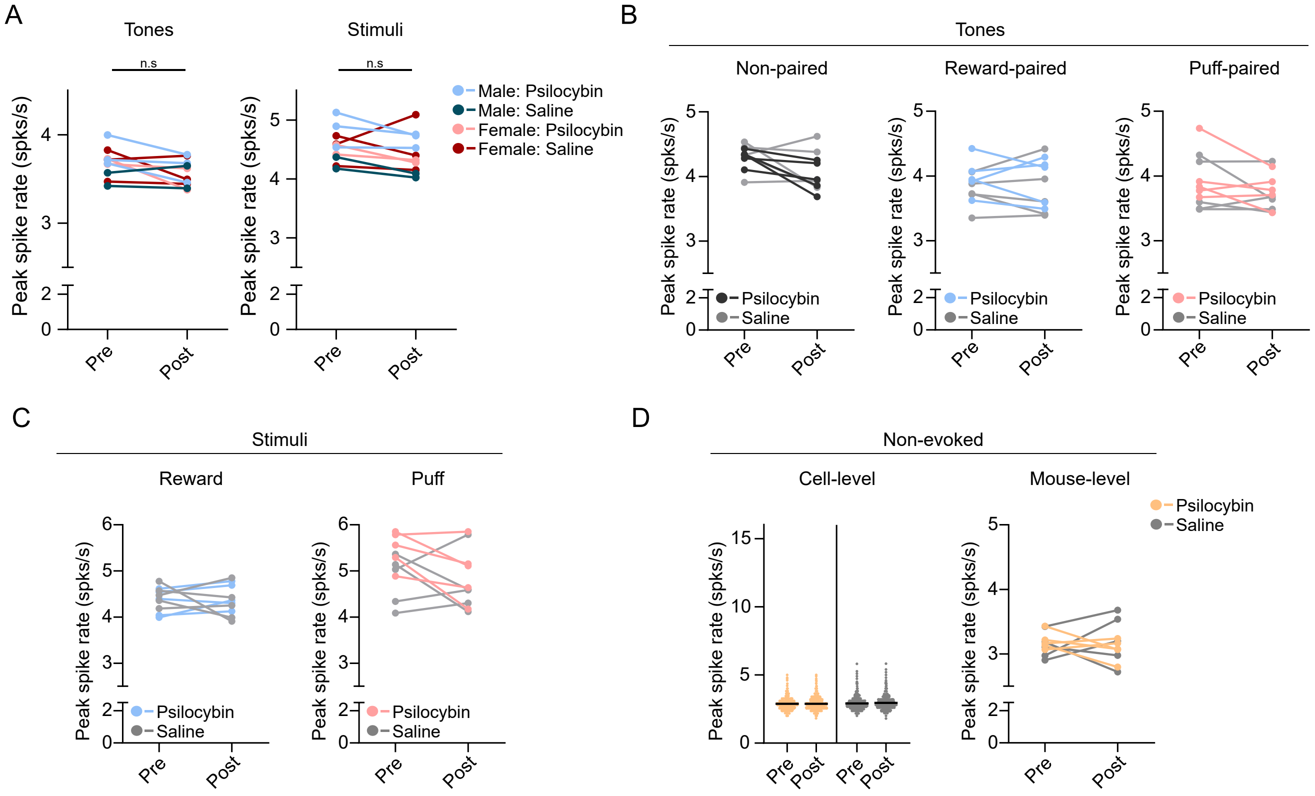


**Supplementary Figure 3 Sex-specific analyses and mouse level data for the multi-valence cohort**

(A) Average peak spike rates for responsive cells by sex for each mouse, pooled across all tones (left) and both stimuli combined (right). All data used mouse-level repeated measures mixed-effects model. n = 5 psilocybin, n = 5 saline mice. Saline = grey, Psilocybin = black (non-paired tone), blue (water-paired tone, reward), pink (puff-paired tone, air puff). (B-D) Mouse-level averages of peak spike rates for tones (B), stimuli (C), and non-evoked trials (D), shown at both the cell-averaged and mouse-averaged levels. Repeated measures mixed-effects model, post-hoc Tukey's test. (D) Each dot = one neuron. n = 5 psilocybin, n = 5 saline mice. Saline = grey, Psilocybin = gold.

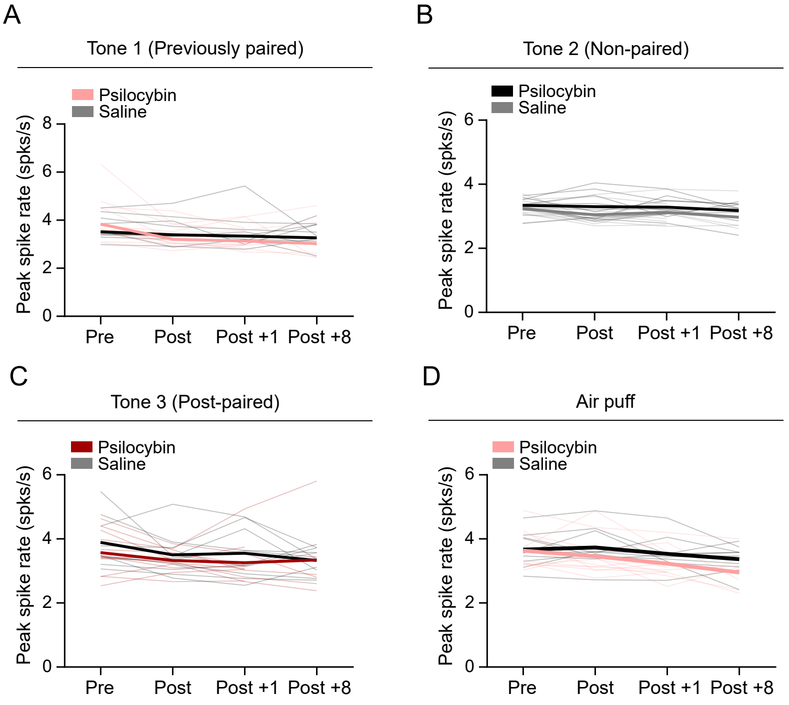


**Supplementary Figure 4. All mouse-level responses in excited neurons**

Average sessional peak spike data from all mice (thin lines) are shown alongside treatment-group means (thick lines) across sessions. neurons. n = 13 psilocybin, n = 10 saline mice per panel. (A) Tone 1 (paired prior to treatment) excited population. Saline = black, Psilocybin = pink. (B) Tone 2 (non-paired) excited population. Saline = black, Psilocybin = light grey. (C) Tone 3 (paired during treatment) excited population. Saline = black, Psilocybin = red. (D) Air-puff responses) excited population. All data: Repeated measures mixed-effects model, post-hoc Tukey's test. Saline = black, Psilocybin = red.

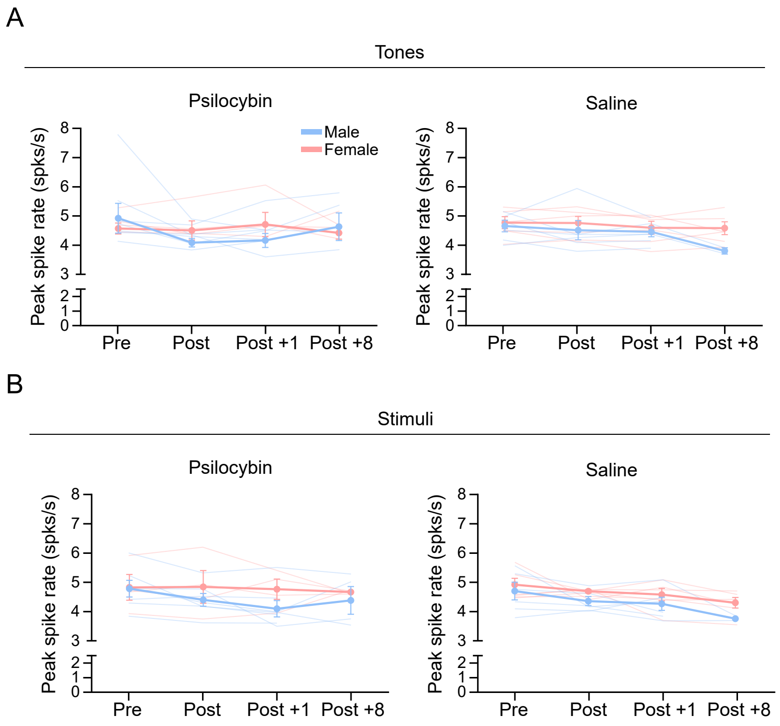


**Supplementary Figure 5. Sex-specific analyses of tone and stimulus responses of aversive only data**

All mice across treatments were split into male and female across session and measures (tones and air puff) to ascertain if any sex differences were present n = 13 psilocybin, n = 10 saline mice per panel. Male = blue, female = pink per panel. (A) Peak spike rates in response to all tones combined, separated by sex and treatment group. Mouse mean ± SEM in thick lines, individual mice in faint thin lines. Repeated measures mixed-effects model, post-hoc Tukey's test. (B) Peak spike rates in response to air puff, separated by sex and treatment group. Mouse mean ± SEM in thick lines, individual mice in faint thin lines. Repeated measures mixed-effects model, post-hoc Tukey's test.

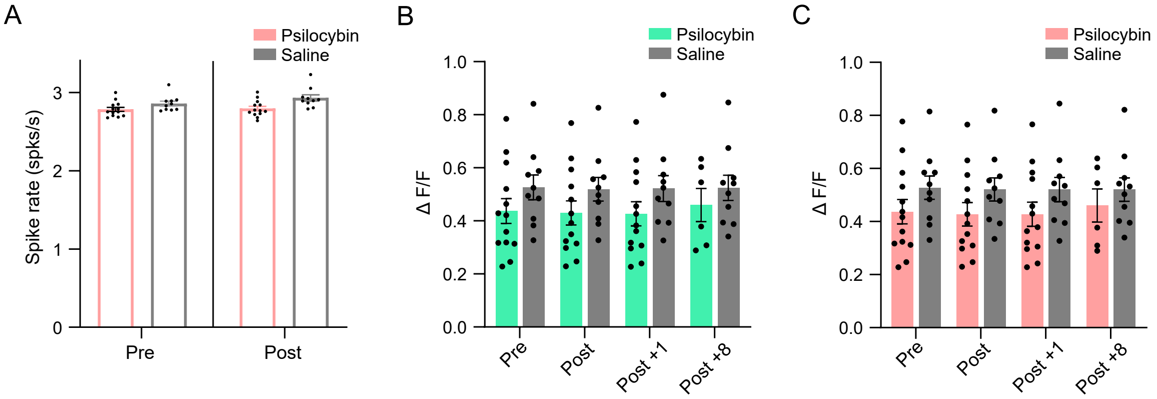


**Supplementary Figure 6. Non-evoked and Inter-trial interval responses**

Non-evoked data was collected as separate blocks of imaging without tones or stimuli prior to and following injections during injection days. Inter-trial interval data was collected in between tones or stimuli during each session. Each dot = one mouse. Bars = mean ± SEM. n = 13 psilocybin, n = 10 saline mice per panel. (A) Non-evoked neuronal activity across all cells before and after injection. Average spike rates were calculated across the 8-minute spontaneous recording blocks. One-way ANOVA. Each dot is one mouse. (B, C) Inter-trial interval (ITI) GCaMP8f ΔF/F measured between tones and air-puff stimuli respectfully. Two-way repeated measures ANOVA. Saline = grey, psilocybin = green (tones) or pink (air puff).

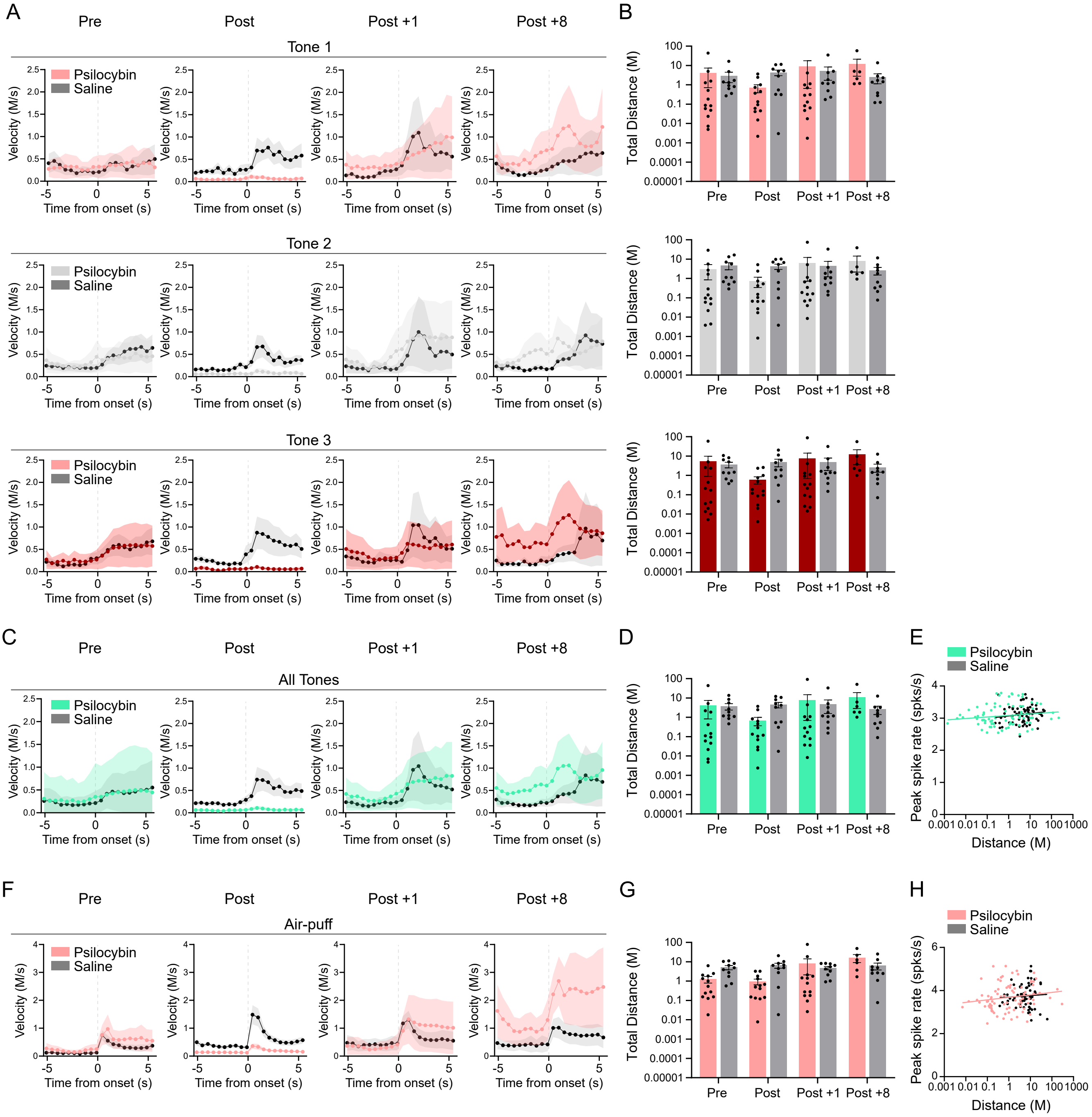


**Supplementary Figure 7. Movement does not account for psilocybin-induced changes in auditory cortical responses**
Movement on a spherical treadmill was continuously recorded and aligned to imaging sessions. n = 13 psilocybin, n = 10 saline mice per panel. (A) Treatment group averages of locomotor activity during tone response windows for each tone across week 2 & 3 sessions (top: previously paired tone 1; middle: non-paired tone 2; bottom: post-paired tone 3). Thick lines and dots = all mouse average, shaded area = SEM. Saline = black, psilocybin = pink, grey, red. (B) Total distance traveled during post-onset each tone response windows across sessions. Repeated measures mixed-effects model, post-hoc Tukey's test. Each dot = one mouse. Bars = mean ± SEM. (top: previously paired tone 1; middle: non-paired tone 2; bottom: post-paired tone 3). Saline = black, psilocybin = pink, grey, red. (C) Treatment group averages of movement during tone response windows for across all tones. Thick lines and dots = all mouse average, shaded area = SEM. Saline = grey, psilocybin = green. (D) Total distance traveled during post-onset all tone response windows across sessions. Repeated measures mixed-effects model, post-hoc Tukey's test. Each dot = one mouse. Bars = mean ± SEM. (E) Correlation between total distance traveled per response window and peak neuronal spike rate across cells during the post-onset tone response windows. Linear regression analysis. Saline = grey, psilocybin = green. Each dot represents one session per mouse. (F) Treatment group averages of movement during air puff response windows. Thick lines and dots = all mouse average, shaded area = SEM. Saline = grey, psilocybin = pink. (G) Total distance traveled during post-onset air puff response windows across sessions. Repeated measures mixed-effects model, post-hoc Tukey's test. Each dot = one mouse. Bars = mean ± SEM. (H) Correlation between total distance traveled per response window and peak neuronal spike rate across cells during the post-onset air puff response windows. Linear regression analysis. Saline = grey, psilocybin = pink. Each dot represents one session per mouse.

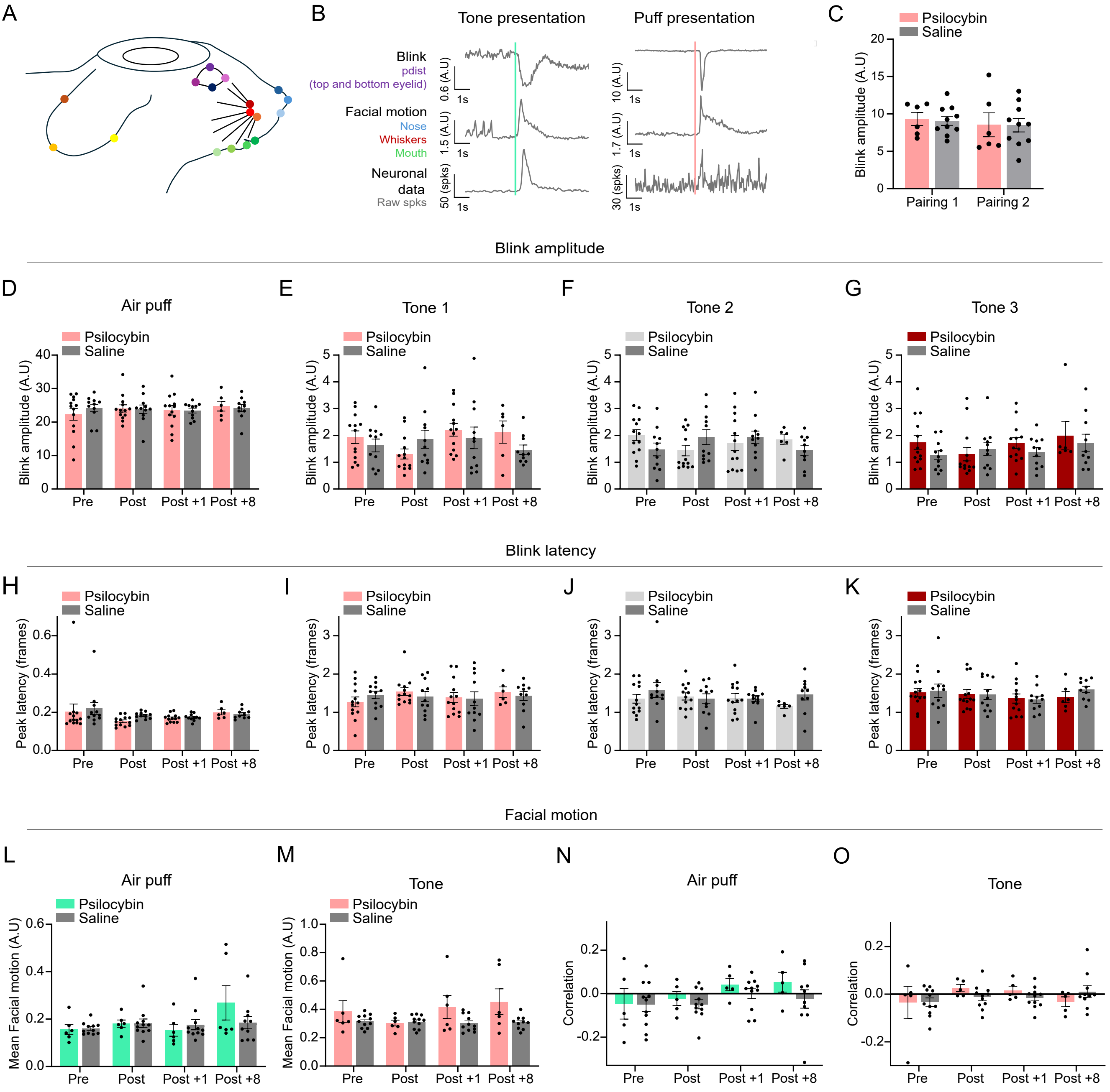


**Supplementary Figure 8. Facial behavior does not account for psilocybin-induced changes in auditory cortical responses**

Videos of facial features and blinks were recorded with an infrared camera and processed using DeepLabCut trained to track facial landmarks (nose, whiskers, mouth, ears) and blink responses (eye corners and eyelids). Repeated measures mixed-effects model [treatment x session x interaction], post-hoc Tukey's test all panels (D-M) unless stated. Each dot = one mouse. Bars = mean ± SEM. (A) Representative image showing DeepLabCut-tracked facial features used for behavioral analysis. (B) Example traces of aligned response windows showing blink amplitude (top-bottom eyelid distance), facial motion (averaged displacement of nose, whisker, and mouth landmarks), and corresponding neuronal raw spikes for the same trial. (C) Peak blink amplitude (within 3 s of tone onset) during pairing sessions in weeks 1 and 2 across treatment groups. Saline = dark grey, psilocybin = pink. n = 6 psilocybin, n = 10 saline mice. (D) Peak blink amplitude in response to air puffs across sessions and treatment groups. Saline = dark grey, psilocybin = pink. (E-G) Peak blink amplitude in response to tones 1, 2 and 3 across sessions and treatment groups. Saline = dark grey, psilocybin = pink, light grey, red. (H) Peak blink latency in response to air puffs across sessions and treatment groups. Saline = dark grey, psilocybin = pink, light grey, red. (I-K) Peak blink latency in response to tone 1, 2 and 3 across sessions and treatment groups. Saline = black, psilocybin = pink. (L, M) Average facial motion in response to all tones (L) and air puffs (M) across treatment groups and sessions. Saline = dark grey, psilocybin = green, pink. n = 6 psilocybin, n = 11 saline mice. (N, O) Correlations between facial motion and peak neuronal spike rates for tone (N) and air-puff (O) responses, shown separately by treatment group and session. Saline = dark grey, psilocybin = green, pink. n = 6 psilocybin, n = 11 saline mice.
