## Supplementary material for "Psilocybin Attenuates Cortical Representations of Aversion in the Mouse Auditory Cortex": Statistics Table

| Figure | Panel | Outcome | Experimental_condition | Test | Test_statistic | p_value | df | p adjustment & notes | N_mice (psy, sal) | N_cells: psy | N_cells: sal |
|---|---|---|---|---|---|---|---|---|---|---|---|
| 1 | E | **Treatment vs session (% of subpopulation)** | **Tone and stimuli response categories** | **Compositional Data Analysis** | **Aitchison Distance** | **p value** | | type III Wald chi-square tests from linear mixed-effects models with Mouse as random effect | 5, 5 | | |
| | | | Aitchison Distance | Pre (Sal vs PSY) | 2.405 | 0.158 | | | | 333 | 521 |
| | | | | Post (Sal vs PSY) | 0.831 | 0.991 | | | | 229 | 498 |
| | | | | **Chi-square test** | **Chi-squared** | **p value** | | | 5, 5 | | |
| | | | V1 (Tone-only vs others) | Group | 0.067 | 0.796 | 1.00 | | | | |
| | | | | Session | 1.683 | 0.195 | 1.00 | | | | |
| | | | | Group:Session | 0.986 | 0.321 | 1.00 | | | | |
| | | | V2 (mixed-balances) | Group | 1.953 | 0.162 | 1.00 | | | | |
| | | | | Session | 0.084 | 0.772 | 1.00 | | | | |
| | | | | Group:Session | 1.461 | 0.227 | 1.00 | | | | |
| | | | V3 (mixed-balances) | Group | 2.635 | 0.105 | 1.00 | | | | |
| | | | | Session | 1.753 | 0.185 | 1.00 | | | | |
| | | | | Group:Session | 0.002 | 0.969 | 1.00 | | | | |
| | | | V4 (mixed-balances) | Group | 0.001 | 0.981 | 1.00 | | | | |
| | | | | Session | 0.053 | 0.819 | 1.00 | | | | |
| | | | | Group:Session | 0.020 | 0.887 | 1.00 | | | | |
| | | | V5 (mixed-balances) | Group | 0.315 | 0.575 | 1.00 | | | | |
| | | | | Session | 1.350 | 0.245 | 1.00 | | | | |
| | | | | Group:Session | 0.369 | 0.543 | 1.00 | | | | |
| | | | V6 (mixed-balances) | Group | 1.809 | 0.179 | 1.00 | | | | |
| | | | | Session | 18.967 | 0.00001 *** | 1.00 | | | | |
| | | | | Group:Session | 1.655 | 0.198 | 1.00 | | | | |
| 1 | H-left | **Treatment vs session (Peak spike rate)** | **Non-paired** | **Repeated-measures mixed model** | **F value** | **p value** | | Mouse and unique cell as random effects | 5, 5 | 171 | 137 |
| | | | | Treatment | 0.110 | 0.741 | 1, 306 | | | | |
| | | | | Session | 18.009 | 2.918e-05 *** | 1, 306 | | | | |
| | | | | Treatment x sesson interaction | 0.148 | 0.700 | 1, 306 | | | | |
| 1 | H-middle | **Treatment vs session (Peak spike rate)** | **Water-paired** | **Repeated-measures mixed model** | **F value** | **p value** | | Mouse and unique cell as random effects | 5, 5 | 234 | 229 |
| | | | | Treatment | 1.192 | 0.307 | 1, 7.81 | | | | |
| | | | | Session | 2.230 | 0.136 | 1, 461 | | | | |
| | | | | Treatment x sesson interaction | 3.182 | 0.075 | 1, 461 | | | | |
| 1 | H-right | **Treatment vs session (Peak spike rate)** | **Puff-paired** | **Repeated-measures mixed model** | **F value** | **p value** | | Mouse and unique cell as random effects | 5, 5 | 227 | 208 |
| | | | | Treatment | 0.264 | 0.621 | 1, 8 | | | | |
| | | | | Session | 14.657 | 0.000148 *** | 1, 433 | | | | |
| | | | | Treatment x sesson interaction | 0.019 | 0.891 | 1, 433 | | | | |
| 1 | I-left | **Treatment vs session (Peak spike rate)** | **Water** | **Repeated-measures mixed model** | **F value** | **p value** | | Mouse and unique cell as random effects | 5, 5 | 119 | 135 |
| | | | | Treatment | 0.002 | 0.969 | 1, 7.49 | | | | |
| | | | | Session | 0.008 | 0.929 | 1, 252 | | | | |
| | | | | Treatment x sesison interaction | 5.746 | 0.01726 * | 1, 252 | | | | |
| | | | | **Post-hoc comparisons: Holm-Bonferroni** | | | | Mouse and unique cell as random effects | | | |
| | | | | Psy: Pre vs Post | -1.583 | 0.115 | 252.00 | | | | |
| | | | | Sal: Pre vs Post | 1.816 | 0.071 | 252.00 | | | | |
| 1 | I-right | **Treatment vs session (Peak spike rate)** | **Air puff** | **Repeated-measures mixed model** | **F value** | **p value** | | Mouse and unique cell as random effects | 5, 5 | 274 | 224 |
| | | | | Treatment | 2.447 | 0.155 | 1, 8.3 | | | | |
| | | | | Session | 10.780 | 0.00100 ** | 1, 496 | | | | |
| | | | | Treatment x sesson interaction | 10.912 | 0.00103 ** | 1, 496 | | | | |
| | | | | **Post-hoc comparisons: Holm-Bonferroni** | | | | Mouse and unique cell as random effects | | | |
| | | | | Psy: Pre vs Post | 4.910 | <0.0001 | 496.00 | | | | |
| | | | | Sal: Pre vs Post | -0.013 | 0.989 | 496.00 | | | | |
| 1 | J | **Session per Treatment (Peak spike rate)** | **Tones: Water-paired/air puff-paired** | **3-way repeated-measures mixed model** | **F value** | **p value** | | Mouse and unique cell as random effects | 5, 5 | 1189 | 1261 |
| | | | | Water-paired:airpuff-paired vs treatment vs session | 0.358 | 0.550 | 1, 4892 | | | | |
| | | | | Linear regression analysis: Psilocybin | 39.710 | <0.0001 | 2, 3561 | | | | |
| | | | | Linear regression analysis: Saline | 2.047 | 0.129 | 2, 3777 | | | | |
| 1 | K | **Session per Treatment (Peak spike rate)** | **Stimuli: Water/air puff** | **3-way repeated-measures mixed model** | **F value** | **p value** | | Mouse and unique cell as random effects | 5, 5 | 1189 | 1261 |
| | | | | Water:airpuff vs treatment vs session | 4.657 | 0.031 * | 1, 4892 | | | | |
| | | | | Linear regression analysis: Psilocybin | 7.151 | 0.0008 ** | 2, 3561 | | | | |
| | | | | Linear regression analysis: Saline | 1.105 | 0.331 | 2, 3777 | | | | |
| 2 | C | **Tone preference (% of subpopulation)** | **Treatment vs session vs prefered tone %** | **Compositional Data Analysis** | **Chi-squared** | **p value** | | Type III Wald chi-square tests from linear mixed-effects models with Mouse as random effec | | 912, 795, 984, 280 | 615, 601, 657, 520 |
| | | | V1 (Tone 1 vs Others) | Session | 0.083 | 0.773 | 1.00 | | 13,10 | | |
| | | | | Tone | 3.371 | 0.338 | 1.00 | | | | |
| | | | | Group × Session | 3.371 | 0.338 | 3.00 | | | | |
| | | | V2 (Tone 2 vs Tone 3) | Session | 0.031 | 0.860 | 1.00 | | 13,10 | | |
| | | | | Tone | 2.448 | 0.485 | 3.00 | | | | |
| | | | | Group × Session | 2.448 | 0.485 | 3.00 | | | | |
| 2 | D | **Treatment vs session (% of population)** | **All tones** | **Repeated-measures mixed model** | **F value** | **p value** | | Mouse and unique cell as random effects | 13,10 | 2450 | 1699 |
| | | | | Treatment | 1.081 | 0.310 | 1,20.861 | | | | |
| | | | | Session | 3.764 | 0.01557 * | 3, 56.616 | | | | |
| | | | | Treatment x sesson interaction | 0.268 | 0.848 | 3, 56.616 | | | | |
| 2 | E | **Treatment vs session (% of population)** | **Tone 1 (Water-paired)** | **Repeated-measures mixed model** | **F value** | **p value** | | Mouse and unique cell as random effects | 13,10 | 294, 240, 286, 81 | 206, 188, 208, 160 |
| | | | | Treatment | 1.997 | 0.172 | 1, 21.216 | | | | |
| | | | | Session | 2.982 | 0.0388 * | 3, 56.616 | | | | |
| | | | | Treatment x sesson interaction | 0.010 | 0.999 | 3, 56.616 | | | | |
| 2 | E | **Treatment vs session (% of population)** | **Tone 2 (Non-paired)** | **Repeated-measures mixed model** | **F value** | **p value** | | Mouse and unique cell as random effects | 13,10 | 382, 334, 464, 129 | 250, 270, 297, 240 |
| | | | | Treatment | 1.860 | 0.187 | 1, 21.821 | | | | |
| | | | | Session | 3.936 | 0.0127 * | 3, 57.546 | | | | |
| | | | | Treatment x sesson interaction | 0.204 | 0.893 | 3, 57.546 | | | | |
| 2 | E | **Treatment vs session (% of population)** | **Tone 3 (Puff-paired)** | **Repeated-measures mixed model** | **F value** | **p value** | | Mouse and unique cell as random effects | 13,10 | 236, 221, 234, 70 | 159, 143, 152, 120 |
| | | | | Treatment | 0.719 | 0.406 | 1, 20.9 | | | | |
| | | | | Session | 1.197 | 0.319 | 3, 56.549 | | | | |
| | | | | Treatment x sesson interaction | 0.094 | 0.963 | 3, 56.549 | | | | |
| 2 | G | **Treatment vs session (Peak spike rate)** | **Tone 1 (Water-paired)** | **Repeated-measures mixed model** | **F value** | **p value** | | Mouse and unique cell as random effects | 13,10 | 294, 240, 286, 81 | 206, 188, 208, 160 |
| | | | | Treatment | 2.575 | 0.124 | 20.71 | | | | |
| | | | | Session | 4.576 | 0.00339 ** | 1425.85 | | | | |
| | | | | Treatment x sesson interaction | 3.659 | 0.0120 * | 1425.85 | | | | |
| | | | | **Post-hoc comparisons: Holm-Bonferroni** | **t-ratio** | **p value** | | | | | |
| | | | | Pre: Psy vs Sal | 0.418 | 0.678 | 39.60 | | | | |
| | | | | Post: Psy vs Sal | -2.246 | 0.0297 * | 44.30 | | | | |
| | | | | Post +1: Psy vs Sal | -1.077 | 0.288 | 39.70 | | | | |
| | | | | Post +8: Psy vs Sal | -2.006 | 0.0478 * | 93.10 | | | | |
| 2 | H-left | **Post session (Cumulative distribution)** | **Tone 1 (Water-paired)** | **Kolmogorov–Smirnov test:** Post | D = 0.275 | 0.0000380 ** | | | 13,10 | 240, 188 | |
| 2 | H-right | **Post +8 session (Cumulative distribution)** | **Tone 1 (Water-paired)** | **Kolmogorov–Smirnov test:** Post +8 | D = 0.278 | 0.00154 ** | | | 13,10 | 81, 160 | |
| 2 | I | **Treatment vs session (Peak spike rate)** | **Tone 2 (Non-paired)** | **Repeated-measures mixed model** | **F value** | **p value** | | Mouse and unique cell as random effects | 13,10 | 382, 334, 464, 129 | 250, 270, 297, 240 |
| | | | | Treatment | 6.792 | 0.0165 * | 21.02 | | | | |
| | | | | Session | 2.752 | 0.0413 * | 2006.73 | | | | |
| | | | | Treatment x sesson interaction | 0.184 | 0.907 | 2006.73 | | | | |
| 2 | I | **Treatment vs session (Normalized to Pre)** | **Tone 2 (Non-paired)** | **Repeated-measures mixed model** | **F value** | **p value** | | Mouse-level | 13,10 | | |
| | | | | Treatment | 0.998 | 0.329 | 1, 21 | | | | |
| | | | | Session | 3.098 | 0.0339 * | 3, 56 | | | | |
| | | | | Session x Treatment | 0.635 | 0.596 | 3, 56 | | | | |
| 2 | J | **Treatment vs session (Peak spike rate)** | **Tone 3 (Puff-paired)** | **Repeated-measures mixed model** | **F value** | **p value** | | Mouse and unique cell as random effects | 13,10 | 236, 221, 234, 70 | 159, 143, 152, 120 |
| | | | | Treatment | 1.725 | 0.202 | 22.52 | | | | |
| | | | | Session | 4.473 | 0.00393 ** | 1225.07 | | | | |
| | | | | Treatment x sesson interaction | 0.906 | 0.438 | 1225.07 | | | | |
| 2 | K | **Treatment vs Post trial stages (Peak spike rate)** | **All tones** | **Repeated-measures mixed model** | **F value** | **p value** | | Geisser-Greenhouse's | 13,10 | | |
| | | | Paired tone | Trial stage | 108.000 | <0.0001 | 1.597, 31.95 | | | | |
| | | | | Group | 2.055 | 0.167 | 1,20 | | | | |
| | | | | Trial stage x Group | 1.267 | 0.293 | 2,40 | | | | |
| | | | Unpaired tones | Trial stage | 44.510 | <0.0001 | 1.914, 40.19 | | | | |
| | | | | Group | 4.288 | 0.051 | 1,20 | | | | |
| | | | | Trial stage x Group | 0.288 | 0.751 | 2,40 | | | | |
| 2 | L-left | **Treatment vs session (Noise correlation)** | **Tone 1 (Water-paired)** | **Repeated-measures mixed model** | **F value** | **p value** | | Mouse level | 11,10 | 2161 | 1699 |
| | | | | Treatment | 0.012 | 0.913 | 1, 20 | | | | |
| | | | | Session | 0.339 | 0.742 | 2.275, 39.43 | | | | |
| | | | | Treatment x sesson interaction | 3.789 | 0.0265 * | 2.275, 39.43 | | | | |
| | | | | **Post-hoc comparisons: Holm-Bonferroni** | **t-ratio** | **p value** | | Mouse level | | | |
| | | | | Pre: Psy vs Sal | 0.054 | >0.9999 | 14.22 | | | | |
| | | | | Post: Psy vs Sal | 1.711 | 0.356 | 17.97 | | | | |
| | | | | Post +1: Psy vs Sal | 2.968 | 0.0393 * | 14.32 | | | | |
| | | | | Post +8: Psy vs Sal | 0.209 | 0.999 | 11.95 | | | | |
| 2 | L-middle | **Treatment vs session (Noise correlation)** | **Tone 2 (Non-paired)** | **Repeated-measures mixed model** | **F value** | **p value** | | Mouse level | 11,10 | 2161 | 1699 |
| | | | | Treatment | 5.727 | 0.417 | 1, 19 | | | | |
| | | | | Session | 0.690 | 0.572 | 2.588, 44.86 | | | | |
| | | | | Treatment x sesison interaction | 1.783 | 0.171 | 2.588, 44.86 | | | | |
| 2 | L-right | **Treatment vs session (Noise correlation)** | **Tone 3 (Puff-paired)** | **Repeated-measures mixed model** | **F value** | **p value** | | Mouse level | 11,10 | 2161 | 1699 |
| | | | | Treatment | 1.087 | 0.309 | 1, 19 | | | | |
| | | | | Session | 3.858 | 0.0225 * | 2.357, 45.58 | | | | |
| | | | | Treatment x sesison interaction | 0.640 | 0.201 | 2.357, 45.58 | | | | |
| 2 | M | **Treatment vs session (Noise correlation)** | **All tones** | **Repeated-measures mixed model** | **F value** | **p value** | | Mouse level | 11,10 | 2161 | 1699 |
| | | | | Treatment | 1.027 | 0.324 | 1, 19 | | | | |
| | | | | Session | 1.033 | 0.376 | 2.411, 41.79 | | | | |
| | | | | Treatment x sesison interaction | 5.682 | 0.00190 ** | 2.411, 41.79 | | | | |
| | | | | **Post-hoc comparisons: Sidak** | **t-ratio** | | | Mouse level | | | |
| | | | | Pre: Psy vs Sal | 0.275 | 0.998 | 18.68 | | | | |
| | | | | Post: Psy vs Sal | 2.893 | 0.0381 * | 18.02 | | | | |
| | | | | Post +1: Psy vs Sal | 1.549 | 0.461 | 14.45 | | | | |
| | | | | Post +8: Psy vs Sal | 0.907 | 0.852 | 13.93 | | | | |
| 3 | C | **Treatment vs session (Peak spike rate)** | **Air puff** | **Repeated-measures mixed model** | **F value** | **p value** | | Mouse level | 13,10 | 414, 446, 368, 86 | 236, 250, 262, 229 |
| | | | | Treatment | 2.931 | 0.102 | 1, 21.14 | | | | |
| | | | | Session | 23.148 | 1.108e-14 *** | 3, 1679.27 | | | | |
| | | | | Treatment x sesson interaction | 3.798 | 0.00990 ** | 3, 1679.27 | | | | |
| | | | | **Post-hoc comparisons: Holm-Bonferroni** | **t-ratio** | **p value** | | Mouse level | 13,10 | | |
| | | | | Pre: Psy vs Sal | -0.005 | 0.996 | 31.40 | | | | |
| | | | | Post: Psy vs Sal | -2.062 | 0.0478 * | 30.40 | | | | |
| | | | | Post +1: Psy vs Sal | -1.667 | 0.106 | 31.00 | | | | |
| | | | | Post +8: Psy vs Sal | -2.131 | 0.0371 * | 62.00 | | | | |
| 3 | D-left | **Post session (Cumulative distribution)** | **Air puff** | **Kolmogorov–Smirnov test:** Post | D = 0.143 | 0.0420 * | | | 13,10 | 446 | 250 |
| 3 | D-right | **Post +8 session (Cumulative distribution)** | **Air puff** | **Kolmogorov–Smirnov test:** Post +8 | D = 0.137 | 0.306 | | | 13,10 | 86 | 229 |
| 3 | E | **Treatment vs Post trial stages (Peak spike rate)** | **Air puff** | **Repeated-measures mixed model** | **F value** | **p value** | | Mouse level | 13,10 | 446 | 250 |
| | | | | Trial stage | 47.450 | < 0.0001 | 1.11, 23.33 | Geisser-Greenhouse's | | | |
| | | | | Treatment | 0.048 | 0.829 | 1,21 | | | | |
| | | | | Trial stage x Treatment | 0.403 | 0.671 | 2, 42 | | | | |
| 3 | F | **Treatment vs session (% of population)** | **Air puff** | **Repeated-measures mixed model** | | | | Mouse level | 13,10 | | |
| | | | | Treatment | 0.418 | 0.525 | 1, 21 | Geisser-Greenhouse's | | 414, 446, 368, 86 | 236, 250, 262, 229 |
| | | | | Session | 3.542 | 0.0280 * | 2.493, 46.54 | | | | |
| | | | | Treatment x sesson interaction | 2.186 | 0.100 | 3, 56 | | | | |
| 3 | G | **Treatment vs session (Noise correlation)** | **Air puff** | **Repeated-measures mixed model** | **F value** | **p value** | | Mouse level | 13,10 | 2450 | 1699 |
| | | | | Treatment | 0.3425 | 0.565 | 1, 20 | | | | |
| | | | | Session | 0.382 | 0.262 | 2.127, 42.53 | | | | |
| | | | | Treatment x sesson interaction | 1.197 | 0.319 | 3,60 | | | | |
| 4 | B | **Treatment vs session (% of subpopulation)** | **Tone and stimuli response categories** | **Compositional Data Analysis** | **Chi-squared** | **p value** | | Type III Wald chi-square tests from linear mixed-effects models with Mouse as random effect | | 930, 951, 968, 291 | 567, 560, 594, 524 |
| | | | V1 (Tone-only vs Others) | Session | 0.108 | 0.742 | 3.00 | | 13,10 | | |
| | | | | Group | 6.905 | 0.075 | 1.00 | | | | |
| | | | | Group × Session | 10.377 | 0.0160 * | 3.00 | | | | |
| | | | V1 (Tone-only vs Others) | **Post-hoc comparisons: Bonferroni** | **t-ratio** | **p value** | | | | | |
| | | | | Pre: Psy vs Sal | 0.270 | 0.188 | 87.10 | | | | |
| | | | | Post: Psy vs Sal | 0.225 | 0.271 | 87.10 | | | | |
| | | | | Post +1: Psy vs Sal | -0.112 | 0.583 | 87.10 | | | | |
| | | | | Post +8: Psy vs Sal | -0.545 | 0.0309 * | 87.10 | | | | |
| | | | V2 (Air puff-only vs Tone & air puff) | Session | 4.423 | 0.0354 * | 3.00 | | 13,10 | | |
| | | | | Group | 9.775 | 0.0205 * | 1.00 | | | | |
| | | | | Group × Session | 1.939 | 0.585 | 3.00 | | | | |
| 4 | C | **Treatment vs session (Peak spike rate)** | **Tone only (Tone block)** | **Repeated-measures mixed model** | **F value** | **p value** | | Mouse and unique cell as random effects | 13,10 | 516, 505, 600, 205 | 331, 310, 332, 295 |
| | | | | Treatment | 1.737 | 0.585 | 1, 21.21 | | | | |
| | | | | Session | 6.205 | 0.000338 *** | 3, 2608.80 | | | | |
| | | | | Treatment x sesson interaction | 0.869 | 0.456 | 3, 2608.80 | | | | |
| 4 | D | **Treatment vs session (Peak spike rate)** | **Tone & air puff (Tone block)** | **Repeated-measures mixed model** | **F value** | **p value** | | Mouse and unique cell as random effects | 13,10 | 175, 135, 164, 31 | 128, 123, 135, 105 |
| | | | | Treatment | 1.737 | 0.378 | 1, 19.76 | | | | |
| | | | | Session | 2.179 | 0.089 | 3, 694.74 | | | | |
| | | | | Treatment x sesson interaction | 0.708 | 0.547 | 3, 694.74 | | | | |
| 4 | E | **Treatment vs session (Peak spike rate)** | **Air puff only (Air puff block)** | **Repeated-measures mixed model** | **F value** | **p value** | | Mouse and unique cell as random effects | 13,10 | 239, 311, 204, 55 | 108, 127, 127, 124 |
| | | | | Treatment | 1.270 | 0.272 | 1, 22.22 | | | | |
| | | | | Session | 9.487 | 3.509e-06 *** | 3, 987.73 | | | | |
| | | | | Treatment x sesson interaction | 4.145 | 0.0062 ** | 3, 987.73 | | | | |
| | | | | **Post-hoc comparisons: Holm-Bonferroni** | **t-ratio** | **p value** | | | | | |
| | | | | Pre: Psy vs Sal | 0.872 | 0.388 | 43.50 | | | | |
| | | | | Post: Psy vs Sal | -0.595 | 0.556 | 37.40 | | | | |
| | | | | Post +1: Psy vs Sal | -1.168 | 0.250 | 40.90 | | | | |
| | | | | Post +8: Psy vs Sal | -2.454 | 0.0161 * | 86.60 | | | | |
| 4 | F | **Treatment vs session (Peak spike rate)** | **Air puff & tone (Air puff block)** | **Repeated-measures mixed model** | **F value** | **p value** | | Mouse and unique cell as random effects | 13,10 | 175, 135, 164, 31 | 128, 123, 135, 105 |
| | | | | Treatment | 2.633 | 0.120 | 1, 21.10 | | | | |
| | | | | Session | 13.868 | 8.518e-09 *** | 3, 710.05 | | | | |
| | | | | Treatment x sesson interaction | 1.075 | 0.359 | 3, 710.05 | | | | |

| Figure | Panel | Outcome | Experimental_condition | Test | Test statistic | p value | df | p adjustment & notes | N mice (psy, sal) | N_cells: psy | N_cells: sal |
|---|---|---|---|---|---|---|---|---|---|---|---|
| **S1** | **B** | **Treatment vs session (SNR)** | **Tones and air puff** | **Repeated-measures mixed model** | **F value** | **p value** | | Geisser-Greenhouse | 13, 10 | | |
| | | | | Treatment | 0.034 | 0.856 | 1,21 | | | | |
| | | | | Session | 1.201 | 0.311 | 1.944, 37.81 | | | | |
| | | | | Treatment x sesson interaction | 0.902 | 0.525 | 9, 175 | | | | |
| **S1** | **C** | **Treatment vs session (Laser Power)** | **Tones and air puff** | **Repeated-measures mixed model** | **F value** | **p value** | | Geisser-Greenhouse | 13, 10 | | |
| **S3** | **B-left** | **Male vs female, vs session** | **All tones: Male vs female** | **Paired t-tests** | **t value** | **p value** | | Mouse level | 5, 5 | | |
| | | | | Pre Psy: Male vs female | 0.747 | 0.509 | 3 | | 3, 2 | | |
| | | | | Pre Sal: Male vs female | -1.209 | 0.313 | 3 | | 3, 2 | | |
| | | | | Post Psy: Male vs female | 0.918 | 0.426 | 3 | | 2, 3 | | |
| | | | | Post Sal: Male vs female | -0.293 | 0.789 | 3 | | 2, 3 | | |
| **S3** | **B-right** | **Male vs female, vs session** | **Stimuli: Male vs female** | **Paired t-tests** | **t value** | **p value** | | Mouse level | 5, 5 | | |
| | | | | Pre Psy: Male vs female | 1.549 | 0.219 | 3 | | 3, 2 | | |
| | | | | Pre Sal: Male vs female | -1.149 | 0.334 | 3 | | 3, 2 | | |
| | | | | Post Psy: Male vs female | 3.881 | 0.0300 * | 3 | | 2, 3 | | |
| | | | | Post Sal: Male vs female | -1.342 | 0.272 | 3 | | 2, 3 | | |
| **S3** | **C-left** | **Treatment vs session (Peak spike rate)** | **Non-paired** | **Repeated-measures mixed model** | **F value** | **p value** | | Mouse level, mouse as random effect | 5, 5 | | |
| | | | | Treatment | 0.182 | 0.680 | 1, 8.5 | | | | |
| | | | | Session | 5.572 | 0.0459 * | 1, 8 | | | | |
| | | | | Treatment x sesson interaction | 0.332 | 0.580 | 1, 8 | | | | |
| **S3** | **C-middle** | **Treatment vs session (Peak spike rate)** | **Water-paired** | **Repeated-measures mixed model** | **F value** | **p value** | | Mouse level, mouse as random effect | 5, 5 | | |
| | | | | Treatment | 0.645 | 0.439 | 1, 11.2 | | | | |
| | | | | Session | 0.087 | 0.776 | 1, 8 | | | | |
| | | | | Treatment x sesson interaction | 0.153 | 0.706 | 1, 8 | | | | |
| **S3** | **C-right** | **Treatment vs session (Peak spike rate)** | **Puff-paired** | **Repeated-measures mixed model** | **F value** | **p value** | | Mouse level, mouse as random effect | 5, 5 | | |
| | | | | Treatment | 0.277 | 0.610 | 1, 10.6 | | | | |
| | | | | Session | 2.536 | 0.150 | 1, 8 | | | | |
| | | | | Treatment x sesson interaction | 0.093 | 0.768 | 1, 8 | | | | |
| **S3** | **D-left** | **Treatment vs session (Peak spike rate)** | **Water** | **Repeated-measures mixed model** | **F value** | **p value** | | Mouse level, mouse as random effect | 5, 5 | | |
| | | | | Treatment | 2.034 | 0.190 | 1, 8.36 | | | | |
| | | | | Session | 0.053 | 0.823 | 1, 8 | | | | |
| | | | | Treatment x sesson interaction | 2.098 | 0.186 | 1, 8 | | | | |
| **S3** | **D-right** | **Treatment vs session (Peak spike rate)** | **Air puff** | **Repeated-measures mixed model** | **F value** | **p value** | | Mouse level, mouse as random effect | 5, 5 | | |
| | | | | Treatment | 1.500 | 0.254 | 1, 8.46 | | | | |
| | | | | Session | 2.341 | 0.165 | 1, 8 | | | | |
| | | | | Treatment x sesson interaction | 0.926 | 0.364 | 1, 8 | | | | |
| **S3** | **E-left** | **Treatment vs session (Peak spike rate)** | **Non-evoked** | **Repeated-measures mixed model** | **F value** | **p value** | | Cell-level, mouse and unique cell as random effects | 5, 5 | 1189 | 1261 |
| | | | | Treatment | 1.842 | 0.176 | 1, 397.4 | | | | |
| | | | | Session | 2.274 | 0.132 | 1, 2448 | | | | |
| | | | | Treatment x sesson interaction | 3.632 | 0.057 | 1, 2448 | | | | |
| **S3** | **E-right** | **Treatment vs session (Peak spike rate)** | **Non-evoked** | **Repeated-measures mixed model** | **F value** | **p value** | | Mouse level, mouse as random effect | 5, 5 | | |
| | | | | Treatment | 1.895 | 0.188 | 1, 16 | | | | |
| | | | | Session | 0.001 | 0.981 | 1, 16 | | | | |
| | | | | Treatment x sesson interaction | 2.147 | 0.162 | 1, 16 | | | | |
| **S4** | **A** | **Treatment vs session (Peak spike rate)** | **Tone 1 (Previously-paired)** | **Repeated-measures mixed model** | **F value** | **p value** | | Mouse level, mouse as random effect | 13, 10 | | |
| | | | | Treatment | 0.723 | 0.405 | 1, 21.3 | | | | |
| | | | | Session | 1.748 | 0.168 | 3,57 | | | | |
| | | | | Treatment x sesson interaction | 0.943 | 0.426 | 3,57 | | | | |
| **S4** | **B** | **Treatment vs session (Peak spike rate)** | **Tone 2 (Non-paired)** | **Repeated-measures mixed model** | **F value** | **p value** | | Mouse level, mouse as random effect | 13, 10 | | |
| | | | | Treatment | 4.508 | 0.0456 * | 1, 21.3 | | | | |
| | | | | Session | 3.153 | 0.0317 * | 3,57 | | | | |
| | | | | Treatment x sesson interaction | 0.161 | 0.922 | 3,57 | | | | |
| | | | | **Post-hoc comparisons: Holm-Bonferroni** | **t value** | **p value** | | | | | |
| | | | | Pre: Psy vs Sal | -1.349 | 0.183 | 53.2 | | | | |
| | | | | Post: Psy vs Sal | -1.602 | 0.115 | 53.2 | | | | |
| | | | | Post +1: Psy vs Sal | -2.061 | 0.0442 * | 53.2 | | | | |
| | | | | Post +8: Psy vs Sal | -1.235 | 0.221 | 67.5 | | | | |
| **S4** | **C** | **Treatment vs session (Peak spike rate)** | **Tone 3 (Post-paired)** | **Repeated-measures mixed model** | **F value** | **p value** | | Mouse level, mouse as random effect | 13, 10 | | |
| | | | | Treatment | 1.942 | 0.178 | 1, 21.7 | | | | |
| | | | | Session | 3.342 | 0.0254 * | 3,57 | | | | |
| | | | | Treatment x sesson interaction | 1.608 | 0.198 | 3,57 | | | | |
| **S4** | **D** | **Treatment vs session (Peak spike rate)** | **Air puff** | **Repeated-measures mixed model** | **F value** | **p value** | | Mouse level, mouse as random effect | 13, 10 | | |
| | | | | Treatment | 2.742 | 0.112 | 1, 21.6 | | | | |
| | | | | Session | 6.243 | 0.00097 *** | 3,57 | | | | |
| | | | | Treatment x sesson interaction | 0.700 | 0.556 | 3,57 | | | | |
| **S5** | **A-left** | **Male vs female, vs session** | **All tones: Male vs female** | **Repeated-measures mixed model** | **F value** | **p value** | | Mouse level | 13, 10 | | |
| | | Psilocybin | | Session | 0.932 | 0.382 | 1.44, 18.72 | | | | |
| | | | | Sex | 0.169 | 0.846 | 2, 14 | | | | |
| | | | | Sex x sesson interaction | 0.909 | 0.498 | 6, 39 | | | | |
| **S5** | **A-right** | **Male vs female, vs session** | **All tones: Male vs female** | **Repeated-measures mixed model** | **F value** | **p value** | | Mouse level | 13, 10 | | |
| | | Saline | | Session | 4.479 | 0.0248 * | 2.1, 18.22 | | | | |
| | | | | Sex | 1.398 | 0.265 | 1,10 | | | | |
| | | | | Sex x sesson interaction | 2.082 | 0.127 | 3,26 | | | | |
| **S5** | **B-left** | **Male vs female, vs session** | **Air puff: Male vs female** | **Repeated-measures mixed model** | **F value** | **p value** | | Mouse level | 13, 10 | | |
| | | Psilocybin | | Session | 3.252 | 0.057 | 1.92, 24.94 | | | | |
| | | | | Sex | 0.767 | 0.483 | 2,14 | | | | |
| | | | | Sex x sesson interaction | 0.609 | 0.722 | 6,39 | | | | |
| **S5** | **B-right** | **Male vs female, vs session** | **Air puff: Male vs female** | **Repeated-measures mixed model** | **F value** | **p value** | | Mouse level | 13, 10 | | |
| | | Saline | | Session | 5.270 | 0.014 * | 2.1, 18.56 | | | | |
| **S6** | **A** | **Treatment vs session (Peak spike rate)** | **Non-evoked** | **Repeated-measures 2-way ANOVA** | **F value** | **p value** | | Geisser-Greenhouse | 13, 10 | | |
| | | | | Treatment | 7.506 | 0.0148 * | 1, 21 | | | | |
| | | | | Session | 3.705 | 0.068 | 1, 21 | | | | |
| | | | | Treatment x sesson interaction | 2.168 | 0.156 | 1, 21 | | | | |
| **S6** | **B** | **Treatment vs session (delta F / F)** | **Tones: Non-evoked** | **Repeated-measures mixed model** | **F value** | **p value** | | Geisser-Greenhouse | 13, 10 | | |
| | | | | Treatment | 1.979 | 0.174 | 1, 21 | | | | |
| | | | | Session | 1.979 | 0.174 | 1.12, 20.8 | | | | |
| | | | | Treatment x sesson interaction | 0.904 | 0.445 | 3, 56 | | | | |
| **S6** | **C** | **Treatment vs session (delta F / F)** | **Air puff: Non-evoked** | **Repeated-measures mixed model** | **F value** | **p value** | | Geisser-Greenhouse | 13, 10 | | |
| | | | | Treatment | 3.235 | 0.168 | 1, 21 | | | | |
| | | | | Session | 2.040 | 0.083 | 1.11, 20.7 | | | | |
| | | | | Treatment x sesson interaction | 0.106 | 0.956 | 3, 56 | | | | |
| **S7** | **B-top** | **Treatment vs session (Sum movement post-onset)** | **Tone 1 (Previously-paired)** | **Repeated-measures mixed model** | **F value** | **p value** | | Geisser-Greenhouse | 13, 10 | | |
| | | | | Treatment | 0.127 | 0.725 | 1, 21 | | | | |
| | | | | Session | 0.876 | 0.335 | 0.8, 15.7 | | | | |
| | | | | Treatment x sesson interaction | 0.784 | 0.508 | 3, 56 | | | | |
| **S7** | **B-middle** | **Treatment vs session (Sum movement post-onset)** | **Tone 2 (Non-paired)** | **Repeated-measures mixed model** | **F value** | **p value** | | Geisser-Greenhouse | 13, 10 | | |
| | | | | Treatment | 0.000 | 0.975 | 1, 21 | | | | |
| | | | | Session | 0.615 | 0.434 | 0.8, 15.7 | | | | |
| | | | | Treatment x sesson interaction | 0.800 | 0.499 | 3, 56 | | | | |
| **S7** | **B-bottom** | **Treatment vs session (Sum movement post-onset)** | **Tone 3 (Post-paired)** | **Repeated-measures mixed model** | **F value** | **p value** | | Geisser-Greenhouse | 13, 10 | | |
| | | | | Treatment | 0.099 | 0.757 | 1, 21 | | | | |
| | | | | Session | 0.623 | 0.413 | 0.8, 15.7 | | | | |
| | | | | Treatment x sesson interaction | 1.083 | 0.364 | 3, 56 | | | | |
| **S7** | **D** | **Treatment vs session (Sum movement post-onset)** | **All tones** | **Repeated-measures mixed model** | **F value** | **p value** | | Geisser-Greenhouse | 13, 10 | | |
| | | | | Treatment | 0.056 | 0.815 | 1, 21 | | | | |
| | | | | Session | 0.719 | 0.379 | 0.79, 14.85 | | | | |
| | | | | Treatment x sesson interaction | 0.892 | 0.451 | 3, 56 | | | | |
| **S7** | **E** | **Treatment vs session (Movement vs Peak spike rate)** | **All tones** | **Pearson correlation** | | | | | | | |
| | | | Psilocybin | r | -0.025 | 0.836 | | | | | |
| | | | | $r^2$ | 0.001 | | | | | | |
| | | | Saline | r | 0.106 | 0.467 | | | | | |
| | | | | $r^2$ | 0.011 | | | | | | |
| **S7** | **G** | **Treatment vs session (Sum movement post-onset)** | **Air puff** | **Repeated-measures mixed model** | **F value** | **p value** | | Geisser-Greenhouse | 13, 10 | | |
| | | | | Treatment | 0.039 | 0.845 | 1, 21 | | | | |
| | | | | Session | 2.247 | 0.093 | 3, 56 | | | | |
| | | | | Treatment x sesson interaction | 2.346 | 0.083 | 3, 56 | | | | |
| **S7** | **H** | **Treatment vs session (Movement vs Peak spike rate)** | **Air puff** | **Pearson correlation** | | | | | | | |
| | | | Psilocybin | r | -0.155 | 0.158 | | | | | |
| | | | | $r^2$ | 0.024 | | | | | | |
| | | | Saline | r | -0.089 | 0.462 | | | | | |
| | | | | $r^2$ | 0.008 | | | | | | |
| **S8** | **C** | **Pairing 1 vs pairing 2 (Blink amplitude)** | **Air puff** | **Repeated-measures mixed model** | **F value** | **p value** | | Geisser-Greenhouse | 6, 10 | | |
| | | | | Treatment | 0.014 | 0.908 | 1, 14 | | | | |
| | | | | Pairing 1 vs pairing 2 | 0.834 | 0.377 | 1, 14 | | | | |
| | | | | Treatment x sesson interaction | 0.016 | 0.900 | 1, 14 | | | | |
| **S8** | **D** | **Treatment vs session (Blink amplitude)** | **Air puff** | **Repeated-measures mixed model** | **F value** | **p value** | | Geisser-Greenhouse | 6, 10 | | |
| | | | | Treatment | 0.218 | 0.645 | 1,22 | | | | |
| | | | | Session | 0.351 | 0.762 | 2.62, 50.65 | | | | |
| | | | | Treatment x sesson interaction | 0.964 | 0.416 | 3, 58 | | | | |
| **S8** | **E** | **Treatment vs session (Blink amplitude)** | **Tone 1 (Previously-paired)** | **Repeated-measures mixed model** | **F value** | **p value** | | Geisser-Greenhouse | 13, 11 | | |
| | | | | Treatment | 0.758 | 0.393 | 1,22 | | | | |
| | | | | Session | 1.218 | 0.311 | 2.613, 50.52 | | | | |
| | | | | Treatment x sesson interaction | 1.970 | 0.129 | 3, 58 | | | | |
| **S8** | **F** | **Treatment vs session (Blink amplitude)** | **Tone 2 (Non-paired)** | **Repeated-measures mixed model** | **F value** | **p value** | | Geisser-Greenhouse | 13, 11 | | |
| | | | | Treatment | 0.122 | 0.731 | 1,22 | | | | |
| | | | | Session | 0.186 | 0.870 | 2.443,47.23 | | | | |
| | | | | Treatment x sesson interaction | 2.741 | 0.051 | 3, 58 | | | | |
| **S8** | **G** | **Treatment vs session (Blink amplitude)** | **Tone 3 (Post-paired)** | **Repeated-measures mixed model** | **F value** | **p value** | | Geisser-Greenhouse | 13, 11 | | |
| | | | | Treatment | 1.025 | 0.322 | 1,22 | | | | |
| | | | | Session | 1.697 | 0.191 | 2.296, 44.39 | | | | |
| | | | | Treatment x sesson interaction | 1.026 | 0.388 | 3, 58 | | | | |
| **S8** | **H** | **Treatment vs session (Blink latency)** | **Air puff** | **Repeated-measures mixed model** | **F value** | **p value** | | Geisser-Greenhouse | 13, 11 | | |
| | | | | Treatment | 0.411 | 0.528 | 1, 22 | | | | |
| | | | | Session | 2.193 | 0.152 | 1.1, 21.7 | | | | |
| | | | | Treatment x sesson interaction | 0.305 | 0.613 | 1.1, 21.7 | | | | |
| **S8** | **I** | **Treatment vs session (Blink latency)** | **Tone 1 (Previously-paired)** | **Repeated-measures mixed model** | **F value** | **p value** | | Geisser-Greenhouse | 13, 11 | | |
| | | | | Treatment | 0.036 | 0.852 | 1, 22 | | | | |
| | | | | Session | 0.493 | 0.679 | 2.8, 55.1 | | | | |
| | | | | Treatment x sesson interaction | 0.655 | 0.576 | 2.8, 55.1 | | | | |
| **S8** | **J** | **Treatment vs session (Blink latency)** | **Tone 2 (Non-paired)** | **Repeated-measures mixed model** | **F value** | **p value** | | Geisser-Greenhouse | 13, 11 | | |
| | | | | Treatment | 0.958 | 0.338 | 1, 22 | | | | |
| | | | | Session | 0.617 | 0.573 | 2.4, 46.5 | | | | |
| | | | | Treatment x sesson interaction | 1.081 | 0.357 | 2.4, 46.5 | | | | |
| **S8** | **K** | **Treatment vs session (Blink latency)** | **Tone 3 (Post-paired)** | **Repeated-measures mixed model** | **F value** | **p value** | | Geisser-Greenhouse | 13, 11 | | |
| | | | | Treatment | 0.300 | 0.585 | 1, 22 | | | | |
| | | | | Session | 0.960 | 0.407 | 2.6, 69.2 | | | | |
| | | | | Treatment x sesson interaction | 0.272 | 0.817 | 2.6, 69.2 | | | | |
| **S8** | **J** | **Treatment vs session (Facial motion)** | **All tones** | **Repeated-measures mixed model** | **F value** | **p value** | | Geisser-Greenhouse | 6, 10 | | |
| | | | | Treatment | 0.301 | 0.592 | 1, 15 | | | | |
| | | | | Session | 3.037 | 0.068 | 1.855, 27.2 | | | | |
| | | | | Treatment x sesson interaction | 1.714 | 0.201 | 1.855, 27.2 | | | | |
| **S8** | **K** | **Treatment vs session (Facial motion)** | **Air puff** | **Repeated-measures mixed model** | **F value** | **p value** | | Geisser-Greenhouse | 6, 10 | | |
| | | | | Treatment | 3.573 | 0.078 | 1, 15 | | | | |
| | | | | Session | 2.005 | 0.146 | 2.259, 33.13 | | | | |
| | | | | Treatment x sesson interaction | 2.613 | 0.063 | 3,44 | | | | |
| **S8** | **L** | **Treatment vs session (Facial motion vs Peak spike rate)** | **All tones** | **Repeated-measures mixed model** | **F value** | **p value** | | Geisser-Greenhouse | 6, 10 | | |
| | | | | Treatment | 1.636 | 0.222 | 2.384, 32.58 | | | | |
| | | | | Session | 1.736 | 0.187 | 1,14 | | | | |
| | | | | Treatment x sesson interaction | 0.349 | 0.745 | 2.384, 32.58 | | | | |
| **S8** | **M** | **Treatment vs session (Facial motion vs Peak spike rate)** | **Air puff** | **Repeated-measures mixed model** | **F value** | **p value** | | Geisser-Greenhouse | 6, 10 | | |
| | | | | Treatment | 0.098 | 0.759 | 2.345, 32.04 | | | | |
| | | | | Session | 1.002 | 0.389 | 1,14 | | | | |
| | | | | Treatment x sesson interaction | 1.033 | 0.377 | 2.345, 32.04 | | | | |
| | | | | Treatment | 0.957 | 0.339 | 1, 21 | | | | |
| | | | | Session | 0.893 | 0.378 | 1.27, 24.6 | | | | |
| | | | | Treatment x sesson interaction | 2.402 | 0.064 | 9, 175 | | | | |
