## Supplementary material for "Psilocybin Attenuates Cortical Representations of Aversion in the Mouse Auditory Cortex": Resources Table

**Key resources table**

| Reagents and Biological Resources | Source | Identifier |
|---|---|---|
| AAV-syn-GCaMP8f (pGP-AAV-syn-jGCaMP8f-WPRE) | Addgene | # 162376 |
| Dexamethasone 21-phosphate disodium | Sigma-Aldrich (or equivalent) | # 2392-39-4 |
| Xylazine | | |
| Psilocybin (Stock (1 mg/ml in saline)) | Cayman | #14041 |
| Artificial cerebrospinal fluid (aCSF) | Prepared in-house | N/A |
| Ophthalmic eye ointment | Dechra | N/A |
| Dental cement, Puralube | Yates Motloid | # 44115 |
| Mounting medium with DAPI | Southern Biotech | # 0100-20 |
| Cyanoacrylate tissue adhesive (Vetbond) | 3M | #1469SB |
| Cyanoacrylate tissue adhesive | Krazy Glue | KG58548R |
| Ultrasound Gel | Aquasonic | # 03-50 |

**Experimental Models: Organisms/Strains**

| Experimental Models | Source | Identifier |
|---|---|---|
| Mouse: *C57BL/6J* | The Jackson Laboratory | Stock #000664 |

| Equipment | Source | Identifier |
|---|---|---|
| Small animal stereotaxic frame | David Kopf Instruments | Model 940 (or equivalent) |
| Two-photon microscope | Bruker | Ultima / Investigator platform |
| Ti:Sapphire laser (Chameleon Vision S) | Coherent | N/A |
| 25× water-immersion objective (NA 1.05) | Olympus | XLPLN25XWMP2 |
| Orbital nosepiece | Bruker | N/A |
| GaAsP photomultiplier tubes | Hamamatsu | N/A |
| Floating Styrofoam ball treadmill and stimuli systems | PhenoSys | JetBall |
| Infrared camera | Basler | acA1300-60gmNIR |
| Infrared LED array (850 nm) | Thorlabs | LIU850A |
| Ultrasonic audio interface | Avisoft Bioacoustics | UltraSoundGate 116H |
| UV-curable optical adhesive (NOA 71) | Norland Products | NOA 71 |
| Portable LED UV Light Source | U-VIX corporation | # UVC-100 |
| Round glass coverslips (3 mm, 4 mm) | Warner Instruments | # 64-0720/# 64-0724 |
| Hamilton syringe 701RN | Hamilton | # CAL 80330 |
| Hamilton Adaptor | Hamilton | # 55750-01 |
| Glass Capillaries | Harvard Apparatus | # 30-0035 |
| Sugi sponge | Questalpha | #18105-04 |

| Software | Source | Identifier |
|---|---|---|
| MATLAB | MathWorks | https://www.mathworks.com |
| ImageJ / Fiji | NIH | https://imagej.nih.gov |
| Prism 8 or 10 | GraphPad Software | https://www.graphpad.com |
| R | R Core Team | https://www.r-project.org |
| Rstudio (2026.01.1) | Posit | https://posit.co |
| Anaconda | Anaconda Inc. | https://www.anaconda.com |
| Suite2p | Pachitariu et al., 2017 | https://github.com/MouseLand/suite2p |
| DeepLabCut | Nath et al., 2019 | https://github.com/DeepLabCut/DeepLabCut |
| Prairie view | Bruker | |
| Phenosys Control and VR | Phenosys | |

| Code | Source | Identifier |
|---|---|---|
| Custom MATLAB analysis scripts | This study | Available upon request |
